# Cell competition promotes metastatic intestinal cancer through a multistage process

**DOI:** 10.1101/2023.09.14.557359

**Authors:** Ana Krotenberg García, Mario Ledesma-Terrón, Joyce Vriend, Merel E van Luyk, Saskia JE Suijkerbuijk

## Abstract

Cell competition plays an instrumental role in quality control during tissue development and homeostasis. Nevertheless, cancer cells can exploit this process for their own proliferative advantage. In our study, we generated mixed murine organoids and microtissues to explore the impact of cell competition on liver metastasis. Unlike competition at the primary site, the initial effect on liver progenitor cells does not involve the induction of apoptosis. Instead, metastatic competition manifests as a multistage process. Initially, liver progenitors undergo compaction, which is followed by cell cycle arrest, ultimately forcing differentiation. Subsequently, the newly differentiated liver cells exhibit reduced cellular fitness, rendering them more susceptible to outcompetion by intestinal cancer cells. Notably, cancer cells leverage different interactions with different epithelial populations in the liver, using them as scaffolds to facilitate their growth. Consequently, tissue-specific mechanisms of cell competition are fundamental in driving metastatic intestinal cancer.

## Introduction

Worldwide, colorectal cancer is third most common cancer in males and second in females (International Agency for Research on Cancer, WHO). Approximately one-third of colorectal cancer patients develop metastases within the first three years after diagnosis^1–3^. The liver is a main site of colorectal cancer metastasis and is responsible for a large part of the colorectal cancer-dependent lethality.^1,3–6^ Liver tissue is formed by several different cell types that all contribute to optimal liver function, such as detoxification, metabolism and bile production.^7^ The two main epithelial cell types are cholangiocytes and hepatocytes. Cholangiocytes form the bile ducts and are an important source of progenitor cells.^8^ During regeneration these cells can differentiate into hepatocytes as a response to chronic liver damage.^9,10^ This process is characterized by the loss of expression of progenitor markers such as SOX9 and LGR5 and gain expression of typical hepatocyte markers like CYP and Albumin.^8^ Most of the liver is formed by hepatocytes, which represent 60% of the cells and accounts for 80% of the volume of the tissue.^11^ Under homeostatic conditions hepatocytes remain in a quiescent state. However, upon acute injury, hepatocytes can switch to a regenerative state which allows reconstitution of lost tissue through cell proliferation or increasing of cell size.^7,12–14^ The colonization of a secondary organ by cancer cells requires cellular interactions with a novel microenvironment. In the liver this is reflected by various histological growth patterns of liver metastases.^15,16^ Each of these patterns display distinct histological characteristics and often correlating to a specific prognosis. In particular, the replacement pattern, which is characterized by the direct interaction of cancer cells with the surrounding liver epithelium is associated with poor patient outcome.^15^ The importance of interactions between liver cancer cells and surrounding liver tissue is further illustrated by the finding that peritumoral hepatocytes depend on YAP signaling to restrain tumor growth.^17^ However, interactions of metastasis and healthy liver tissue are still poorly understood.

Cell competition is a vital mechanism that drives the continuous selection of cells based on their relative fitness. It serves as a mechanism for quality control during both development and adult tissue homeostasis by elimination of unfit cells.^18^ For example, fluctuations in the expression levels of the transcription factor Myc play a crucial role in determining the survival of cells in the mouse epiblast and developing heart.^19,20^ This mechanism ensures that only the fittest cells constitute these tissues. Additionally, cell competition is employed to remove early malignant cells, characterized by oncogenic Ras expression or a p53 mutation, from epithelial tissues like the pancreas and intestine.^21,22^ It is important to note that cancer cells can exploit this process of cell competition to promote tumor growth.^18,23^ A significant body of research has demonstrated the substantial contribution of cell competition to various stages of primary colorectal cancer. During tumor initiation, APC mutant stem cells gain a crucial competitive advantage over wild-type stem cells through the secretion of the WNT antagonist NOTUM.^24,25^ Additionally, the expression of oncogenic variants of KRas and PI3K leads to a similar competitive advantage.^26^ At later stages of tumorigenesis, we have shown that intestinal cancer cells actively eliminate surrounding healthy epithelial tissue in both the Drosophila adult midgut and murine organoids.^27,28^ In the latter, wild-type cells are eliminated in a JNK dependent manner and activate a fetal-like state, an injury-response that is often activated in damaged intestinal epithelia.^29,30^ Importantly, the cancer population directly benefits from these competitive interactions by boosting their proliferation rate.

To understand the influence of surrounding tissue on growth of colorectal cancer liver metastasis it is crucial to identify the interactions between cancer cells and the epithelial cell populations. Here, we show that intestinal cancer cells outcompete wild-type liver cells in murine organoids and microtissues. When facing progenitor cells, cancer cells induce a cell cycle arrest and subsequent loss of the wild-type progenitor state. The interaction of cancer cells with differentiated, hepatocyte-like cells, causes a reduced cellular fitness and results in rapid outcompetion. Importantly, the cancer population uses wild-type tissue as a scaffold for tumor growth and benefits from these interactions through increased expansion.

## Results

### Cancer cells outcompete wild-type liver cells

To investigate the role of cell competition in liver metastasis we adapted our previously developed 3D mixed organoid model.^31^ For this, membrane-bound tdTomato-labeled wild-type liver cholangiocyte organoids were derived from liver tissue isolated from mTmG transgenic mice as described previously.^32^ These cultures are formed by liver progenitor cells that can be differentiated into functional hepatocytes.^8^ In addition, Dendra2-labeled small intestinal cancer organoids were derived from Villin-CreER^T2^ Apc^fl/fl^ Kras^G12D/WT^ Trp53^fl/R172H^ transgenic mice.^27,33^ To study cellular interactions during liver metastasis, clumps of wild-type liver and small intestinal cancer cells were aggregated to enforce formation of single mixed organoids (Figure 1A).

**Figure 1:**
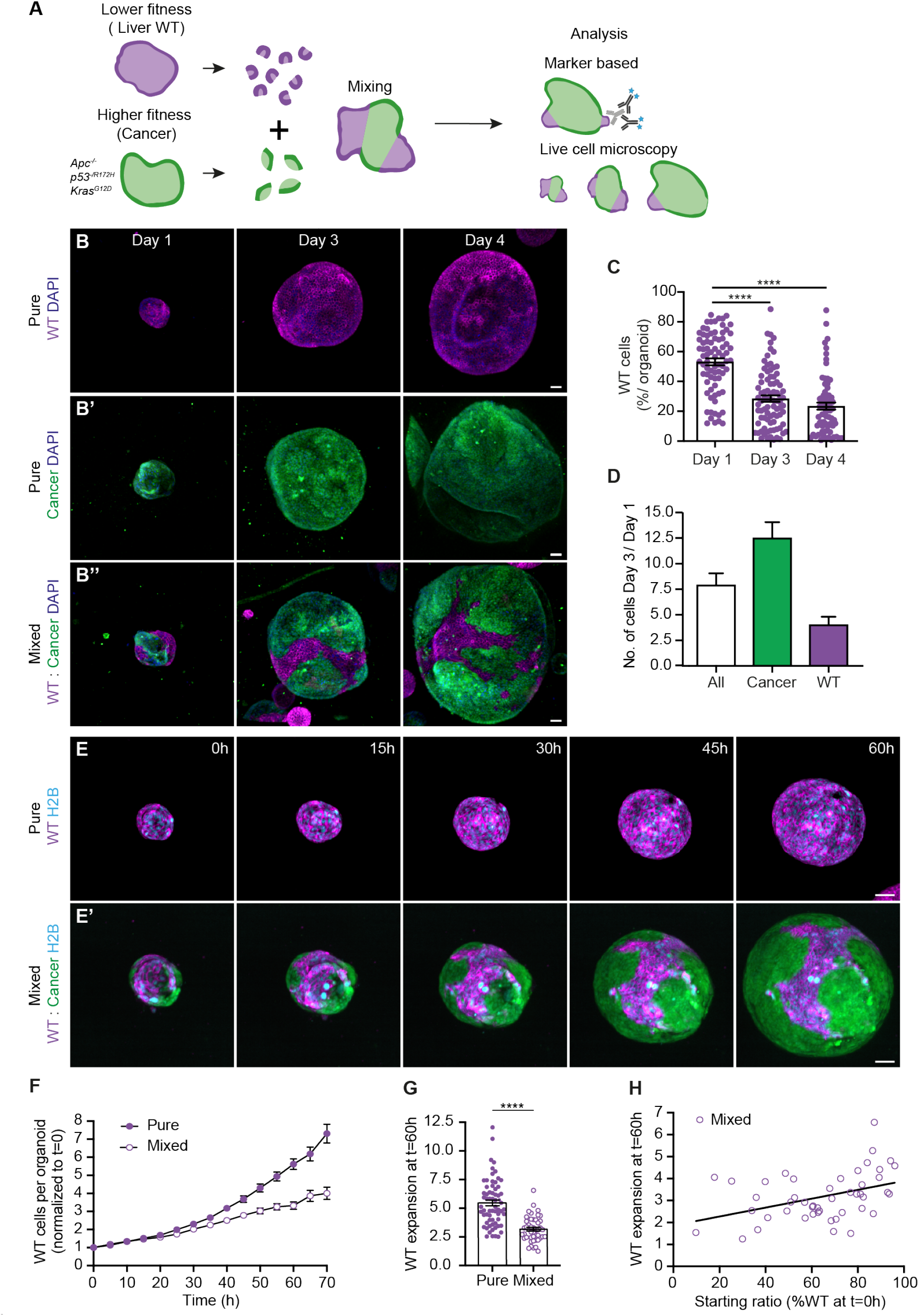
Cancer cells outcompete wild-type liver cells. (A) Schematic depiction of a 3D model for cell competition in murine liver and intestinal cancer organoids. (B) Representative maximum projections of 3D-confocal images of pure WT (B), cancer (B’) and mixed (B’’) organoids fixed at Day 1, 3 and 4 after plating and stained with DAPI (blue). (C) The percentage of WT nuclei contributing to mixed organoids 1, 3 and 4 days after mixing is shown; each dot represents one organoid (mean ± SEM; Kruskal-Wallis test; p<0.0001; n=78, 84 and 69 organoids). (D) Displays the absolute number of cells in organoids shown in (B); the ratio of ‘day 3’ over ‘day 1’ of all (white), cancer (green), and wild-type (magenta) cells are plotted (mean ± SEM; Ordinary one-way ANOVA). (E-H) Analysis of WT cells in pure and mixed H2B-Cerulean3 expressing organoids by live imaging. (E) Representative maximum projections of 3D-confocal images of time-lapse series of WT pure (E) and mixed (E’) organoids. (F) Quantification of the number of WT nuclei in pure and mixed conditions per organoid normalized to the start of the time-lapse (mean ± SEM; paired t-test, two-tailed; p<0.0052; n= 65 and 47 organoids). (G) Quantification of WT expansion, the ratio of the number of WT nuclei at t=60h over t=0h is plotted; each dot represents one organoid (mean ± SEM; Mann-Whitney test, two-tailed; p<0.0001; n= 65 and 47 organoids). (H) Shows the WT expansion at t=60h plotted against the initial percentage of WT cells per mixed organoid (Simple linear regression; R^2^=0.1639; p=0.0048, n=47 organoids). Scale bars represent 50μm. See also Figure S1 and Video S1.

First, we followed the cellular behavior in organoids fixed on day 1, 3 and 4 after plating. Both wild-type and cancer populations increased over a period of four days in pure organoids (Figure 1B). However, the number of wild-type cells contributing to mixed organoids gradually decreased at day 3 (±30%) and 4 (±20%) compared to day 1 (±55%) after mixing (Figure 1C). Furthermore, the total number of cells per organoid showed a 7.5-fold expansion at day 3 compared to day 1, while the absolute number of wild-type cells only increased 2.5 times and the cancer population was responsible for most of the overall expansion (Figure 1D). This indicates that, even though wild-type cells can proliferate, they are outcompeted in mixed organoids.

To gain insight into the dynamics of this competitive behavior, individual nuclei were tracked based on expression of Histone H2B-Cerulean3 using time-lapse microscopy. Pure wild-type organoids showed exponential growth with a doubling time of approximately 24 hours (Figures 1E, 1F and S1A; Video S1). However, the growth rate of wild-type cells was dramatically increased to 33 hours in the presence of cancer cells (Figures 1E, F and S1A; Video S1). Importantly, towards the end of the experiment the average expansion of the wild-type population was reduced from 5.5 times in pure conditions to 3.2 times in mixed organoids (Figure 1G), confirming the outcompetition of wild-type cells. Interestingly, we observed a correlation between the wild-type expansion rate and the percentage of wild-type cells that contributed to the mixed organoid at the start of the experiment. Mixed organoids with a higher percentage of wild-type cells showed an expansion rate that approached pure conditions, while those with a low wild-type contribution also displayed little expansion (Figure 1H). This indicates that the initial number of wild-type cells determines the speed of competition. Together, these data show that wild-type liver cells are outcompeted by cancer cells in mixed organoids.

### Increased expansion of competing cancer

Next, we questioned whether competition could also alter cancer cell behavior. Therefore, we went back to time-lapse microscopy of H2B-Cerulean3 expressing cells (Figure 2A and Video S2). Tracking of the absolute number of cells for up to 70 hours showed that, even though cancer cells proliferated more than wild-type cells in pure organoids (Figures 1F, 2B and S2), the proliferation rate of cancer cells was further increased in mixed organoids (Figures 2B and S2). Furthermore, the average expansion of cancer cells increased from 10.4 in pure to 16.7 times in mixed organoids towards the end of the experiment (Figure 2C). Interestingly, the expansion of cancer cells was inversely correlated to the percentage of cancer cells in the mixed organoid (Figure 2D), suggesting that cancer cells benefit from the presence of relative higher number of wild-type cells. Thus, these data show that wild-type cells provide a growth supportive role that benefits cancer cell expansion.

**Figure 2:**
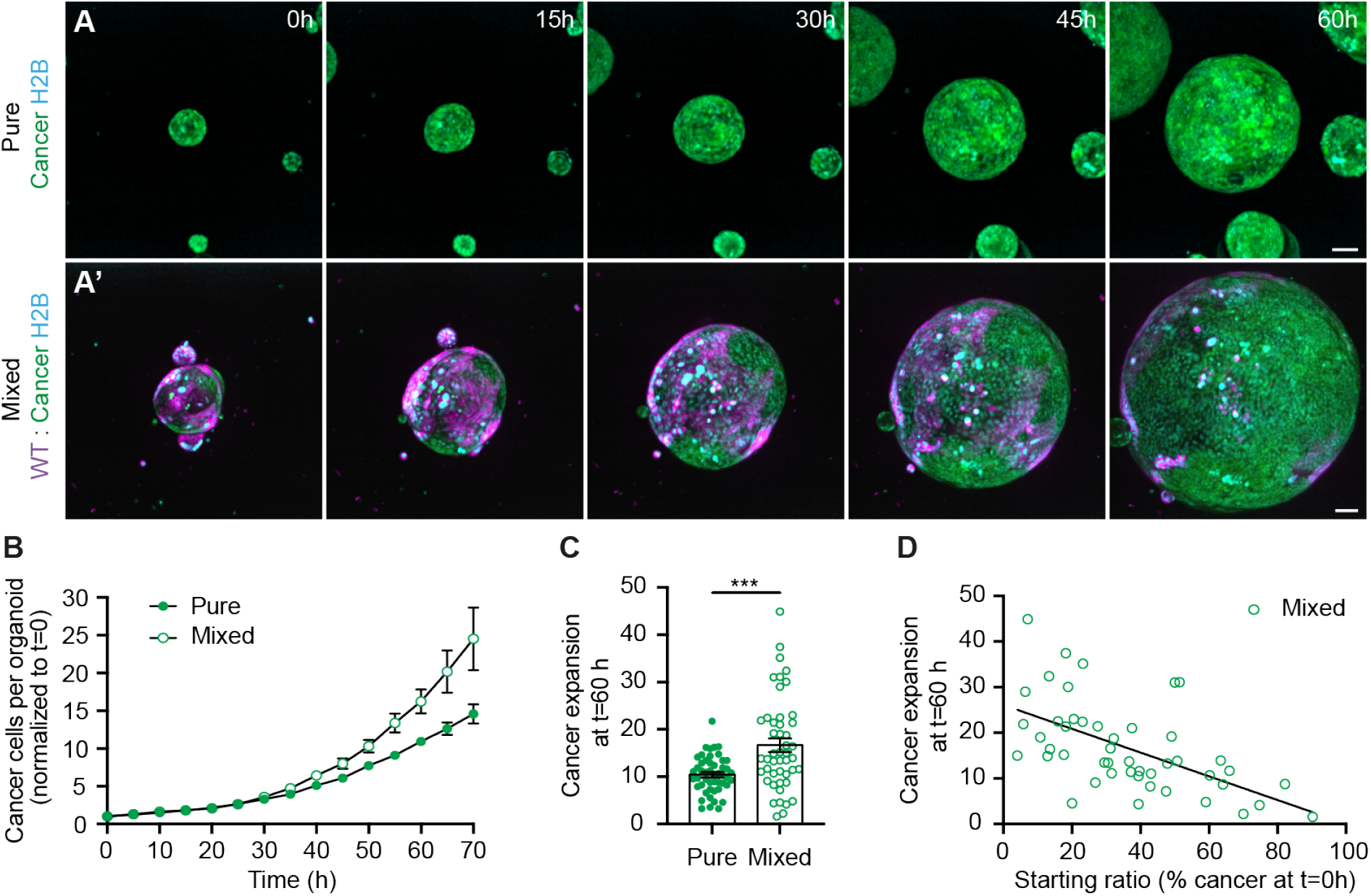
Increased expansion of competing cancer. (A) Representative maximum projections of 3D-confocal images of time-lapse series of H2B-Cerulean3 expressing pure cancer (A) and mixed (A’) organoids. (B) Quantification of the number of cancer nuclei in pure and mixed conditions per organoid normalized to the start of the time-lapse (mean ± SEM; paired t-test, two-tailed; p<0.014; n= 49 and 47 organoids). (C) Quantification of cancer expansion, the ratio of the number of cancer nuclei at t=60h over t=0h is plotted; each dot represents one organoid (mean ± SEM; Mann-Whitney test, two-tailed; p<0.0004; n= 49 and 47 organoids). (D) Shows the cancer expansion at t=60h plotted against the initial percentage of cancer cells per mixed organoid (Simple linear regression; R^2^=0.3229; p<0.0001, n=47 organoids). Scale bars represent 50μm. See also Figure S2 and Video S2.

### Cancer induces compaction and cell cycle arrest of wild-type liver cells

So far, we showed a reduced expansion of the wild-type population in mixed organoids. To further understand the impact of competition on wild-type cells we closely examined interactions between the two cell populations. Two lines of evidence suggest that forces generated by cancer cells induce a morphological change in wild-type cells: 1) The nuclear shape of wild-type cells in mixed organoids appeared more elongated compared to wild-type nuclei in pure organoids (Figure 3A). 2) Measurement of the average distance between the five closest neighbors revealed a decreased inter-nuclear distance of competing wild-type cells (Figures 3B and C). Of note, competition did not affect the inter-nuclear distance of cancer cells in the same organoids (Figures 3B and S3A), indicating that these effects are not a consequence of the overall morphology of the tissue. Together, these data show that competition induces compaction of wild-type cells.

**Figure 3:**
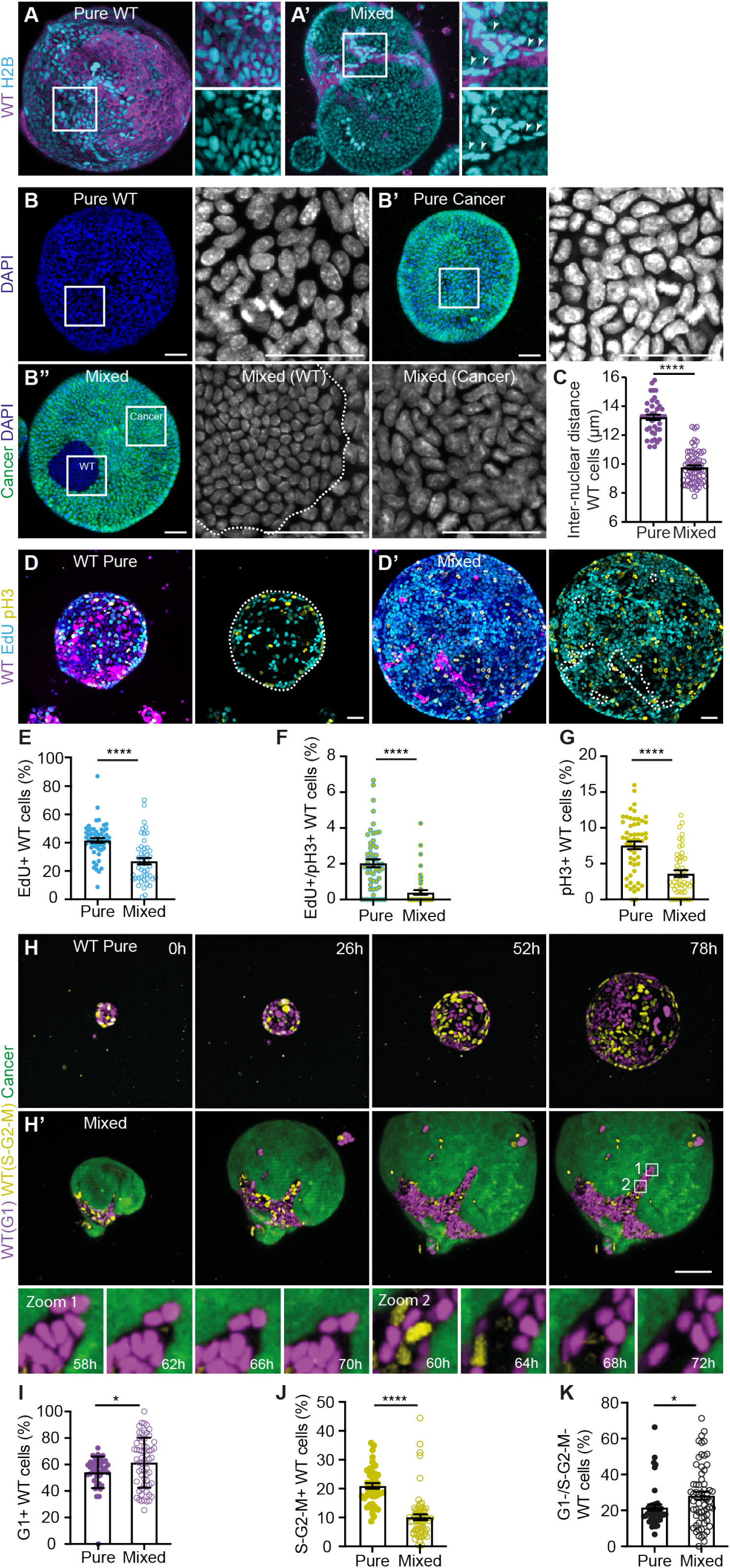
Cancer induces compaction and cell cycle arrest of WT liver cells. (A) Representative maximum projections of 3D-confocal images of pure WT (A) and mixed (A’) organoids. The insets display a 2x magnification of the area in the white box. Nuclei are visualized by expression of H2B-Cerulean3 (cyan). (B) Representative maximum projections of 3D-confocal images of pure WT (B), pure cancer (B’) and mixed (B’’) organoids. The insets display a 2.3x magnification of the area in the white box and white dotted line outlines the wild-type population. Nuclei are visualized with DAPI (grey). (C) Quantification of the inter-nuclear distance of WT cells in pure and mixed organoids; each dot represents one organoid (mean ± SEM; Ordinary one-way ANOVA, Sidak’s multiple comparisons test; p<0.0001; n= 41 and 64 organoids). (D) Representative maximum projections of 3D-confocal images of pure WT (D) and mixed (D’) organoids. Cells in S phase are labeled with EdU (cyan), and mitotic cells are marked by pH3 (yellow); cells that progressed from S phase to mitosis within 1.5h are double positive; the white dotted line outlines the wild-type population. The percentage of WT cells in S phase (E), that progressed from S phase to mitosis (F) and in mitosis (G) pure WT and mixed organoids are plotted; each dot represents one organoid (mean ± SEM; Ordinary one-way ANOVA, Sidak’s multiple comparisons test; p<0.0001; n= 55 and 47 organoids). (H) Representative maximum projections of 3D-confocal images of time-lapse series of pure WT (H) and mixed (H’) organoids. Cell cycle phase of WT cells is visualized by expression of hCDT1-mCherry (G1, magenta) and hGeminin-mVenus (S-G2-M, yellow). The insets display a 7.5x magnification of the area in the white box. (I-K) Quantification of WT cell proliferation in pure and mixed organoids. The percentage of WT cells in G1 (I), S-G2-M (J) and negative (K) are plotted; each dot represents one organoid (mean ± SEM; unpaired t test, two-tailed; p<0.0275, I; p<0.0001, J; p<0.0318, K; n=43 and 62 organoids). Scale bars represent 50μm. See also Figure S3 and S3.

Cell compaction and tissue crowding have previously been described to cause active elimination of less fit cells.^34,35^ However, in the time-lapse movies of mixed organoids we did not find evidence for increased apoptosis or extrusion of wild-type cells. This suggests that the lower expansion rate of competing wild-type cells is instead caused by an effect on cell proliferation. Therefore, we first used two complementary markers of cell proliferation to study the impact of competition on different phases of the cell cycle; DNA replication was visualized by incorporation of the thymidine analogue 5-Ethynyl-2′-deoxyuridine (EdU) and phosphorylation of Histone H3-Ser10 (pH3) was used to recognize mitotic chromatin. The combination of both markers allowed us to identify cells in S phase (EdU+), cells proceeding from S phase to mitosis (EdU+/pH3+) and cells in mitosis (pH3+). Importantly, for all three phases fewer wild-type cells were found in mixed organoids (Figures 3D-G), indicating that competition induces a decrease in wild-type proliferation throughout different phases of the cell cycle.

Next, as cell proliferation is a highly dynamic process, we aimed to visualize cell cycle progression in real-time. For this, wild-type organoids were derived from Fluorescent Ubiquitylation-based Cell Cycle Indicator-2 (FUCCI2) transgenic mice.^36^ This allowed us to follow progression of wild-type cells from the G1 phase (based on hCDT1-mCherry) through S-G2-M phases (based on hGeminin-mVenus) of the cell cycle. Nuclei in pure wild-type organoids showed an alternating expression of hCDT1-mCherry and hGeminin-mVenus throughout the experiment (Figure 3H and Video S3), confirming continued active cell cycle progression of these cells. In contrast, where initially cycling wild-type cells were detected in mixed organoids, this population rapidly decreased to almost absence towards the end of the experiment. Instead, there was an accumulation of the number of hCDT1-mCherry positive wild-type cells at later time-points (Figure 3H and Video S3). These finding were confirmed by quantification of expression of the FUCCI2 reporters in pure and mixed organoids fixed three days after plating (Figure S3D), which showed an increase of G1 and decrease of S-G2-M wild-type cells during competition (Figured 3I and 3J). Interestingly, we observed that a subpopulation of wild-type cells lost both FUCCI2 markers (Figures 3H insets and 3K). These cells arrested after degradation of hGeminin while CDT1 was not yet expressed and were in a putative G0 state. In addition, a similar growth arrested population was detected in the previously described cell proliferation experiment, where we found an increased percentage of wild-type cells that was negative for both EdU and pH3 in mixed organoids (Figure S3B). Even after extending the EdU pulse to 24 hours, ±30% of the competing wild-type population was EdU-/pH3-(Figure S3C), indicating a complete stop in the proliferation of a large population of competing wild-type cells. Together, these data show that cancer cells outcompete wild-type cells through compaction and a subsequent cell cycle arrest.

### Increased competition through loss of WT progenitor state

A cell cycle arrest is typically linked to a high level of cell differentiation.^37^ In addition, several types of cell competition are driven by forced differentiation of stem or progenitor cells. For example, in both the fly and mouse intestine increased differentiation causes loss of less fit cells.^24,25,38^ Furthermore, stem cell displacement from the niche drives cell selection by forced differentiation in the mouse epidermis.^39,40^ Therefore, we next wondered whether the cell cycle arrest of competing wild-type cells is correlated to an altered differentiation state. In adult liver tissue, progenitor cells can trans-differentiate into hepatocytes after chronic liver damage in a process known as ductal reaction and this can be mimicked in organoid cultures.^8,41^ SOX9 is one of the main markers of the liver progenitor state and its expression strongly decreases during differentiation.^42–44^ Therefore, we next focused on expression of this transcription factor. As expected, high nuclear SOX9 expression was found in pure liver progenitor cultures (Figures 4A and S4). Similarly, cancer cells showed heterogeneously high expression of SOX9, which was irrespective of the presence of wild-type cells (Figures 4A and S4). In contrast, competing wild-type cells showed a major reduction in average SOX9 expression (Figures 4A, 4B and S4). Furthermore, the percentage of wild-type cells with high SOX9 expression was severely reduced in mixed organoids (Figure 4C), indicating that fewer progenitor cells remained in the competing wild-type population. On the other hand, the percentage of wild-type cells with low SOX9 expression was strongly increased (Figure 4D), suggesting that competition induces differentiation in most wild-type cells. Interestingly, we observed that organoids with fewer remaining wild-type cells also had more cells with low SOX9 expression (Figure 4E). Together, this suggests that more cancer cells can inflict a stronger response and that the strength of competition is determined by the relative contribution of cell populations.

**Figure 4:**
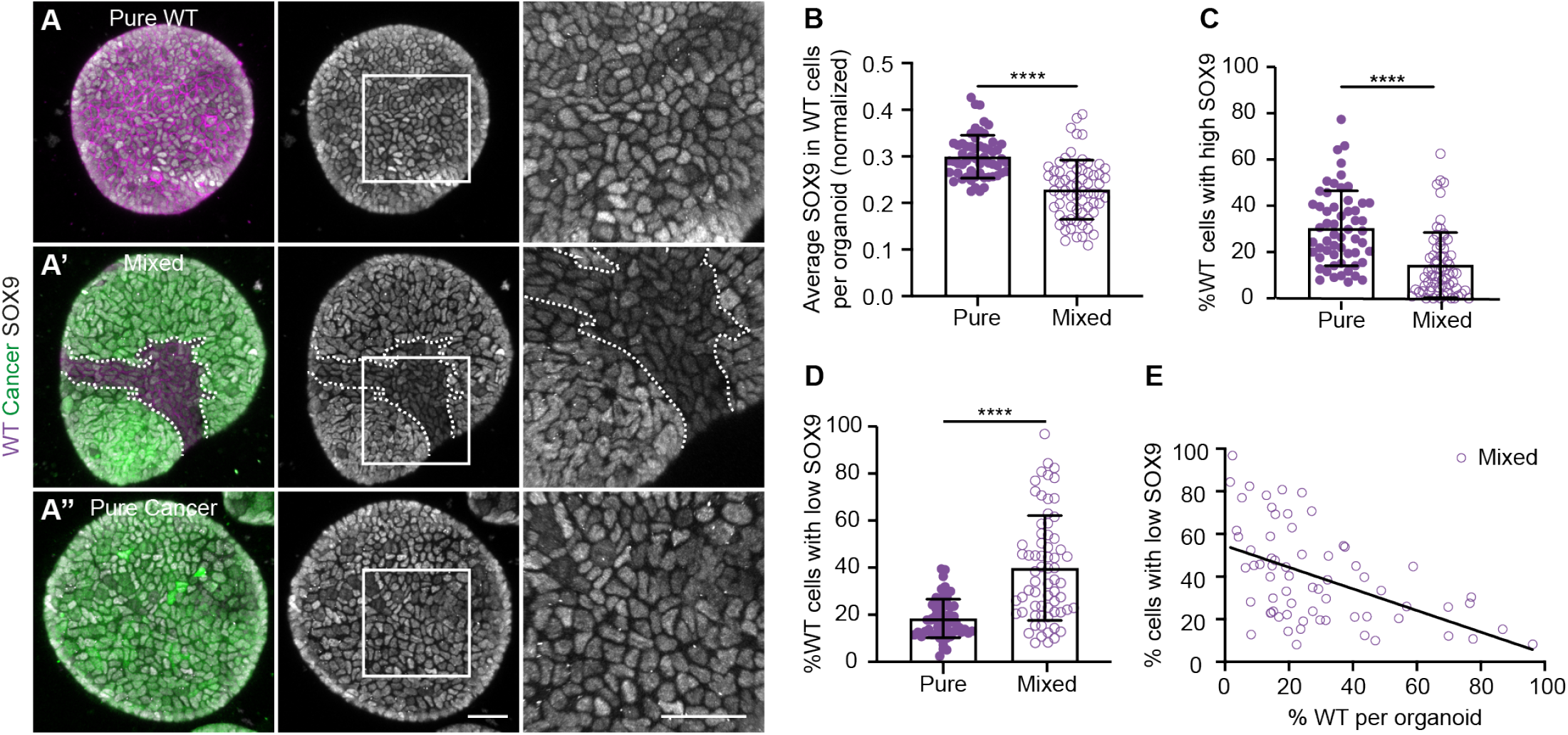
Increased competition through loss of WT progenitor state. (A) Representative maximum projections of 3D-confocal images of WT pure (A), mixed (A’) and cancer pure (A’’) organoids fixed 3 days after plating. The organoids were stained for SOX9 (grey). The insets display a 2.5x magnification of the area in the white box and the white dotted line outlines the wild-type population. (B-E) SOX9 expression in WT cells. (B) Displays the average SOX9 intensity in WT cells in pure and mixed organoids; each dot represents one organoid (mean ± SEM; Mann-Whitney test, two-tailed; p<0.0001; n= 57 and 67 organoids). The percentage of WT cells with high (C) and low (D) SOX9 expression is plotted and each dot represents one organoid (mean ± SEM; Kruskal-Wallis test; p<0.0001; n= 57 and 67 organoids). (E) Shows the percentage of WT cells with low SOX9 levels plotted against the percentage of WT cells per organoid (Simple linear regression; R^2^=0.2511; p<0.0001, n=67 organoids). Scale bars represent 50μm. See also Figure S4.

Thus, cell competition driven by intestinal cancer cells induces a loss of the progenitor state of wild-type liver cells.

### Differentiated WT cells are effectively outcompeted

So far, our findings suggest that competition causes a cell cycle arrest and loss of progenitor state in wild-type cells. We next aimed to analyze the effect of differentiation on the outcome of competition. For this, wild-type liver progenitor cells were exposed to a well-characterized 15 days differentiation protocol (Figure 5A). This generated differentiated hepatocyte-like organoids that lack expression of SOX9 (Figure 5B). Furthermore, in contrast to wild-type progenitors, which showed a 4.5-fold expansion over de course of two days (Figures 5C and S5B), differentiated cells were non-proliferative (Figures 5C and 5D). This was confirmed using time-lapse microscopy (Figures S5A and Video S4). Importantly, the reduced proliferation was not caused by a general cytostatic effect of the differentiation medium as expansion of pure cancer organoids was not affected (Figures S5C-G and Video S5).

**Figure 5:**
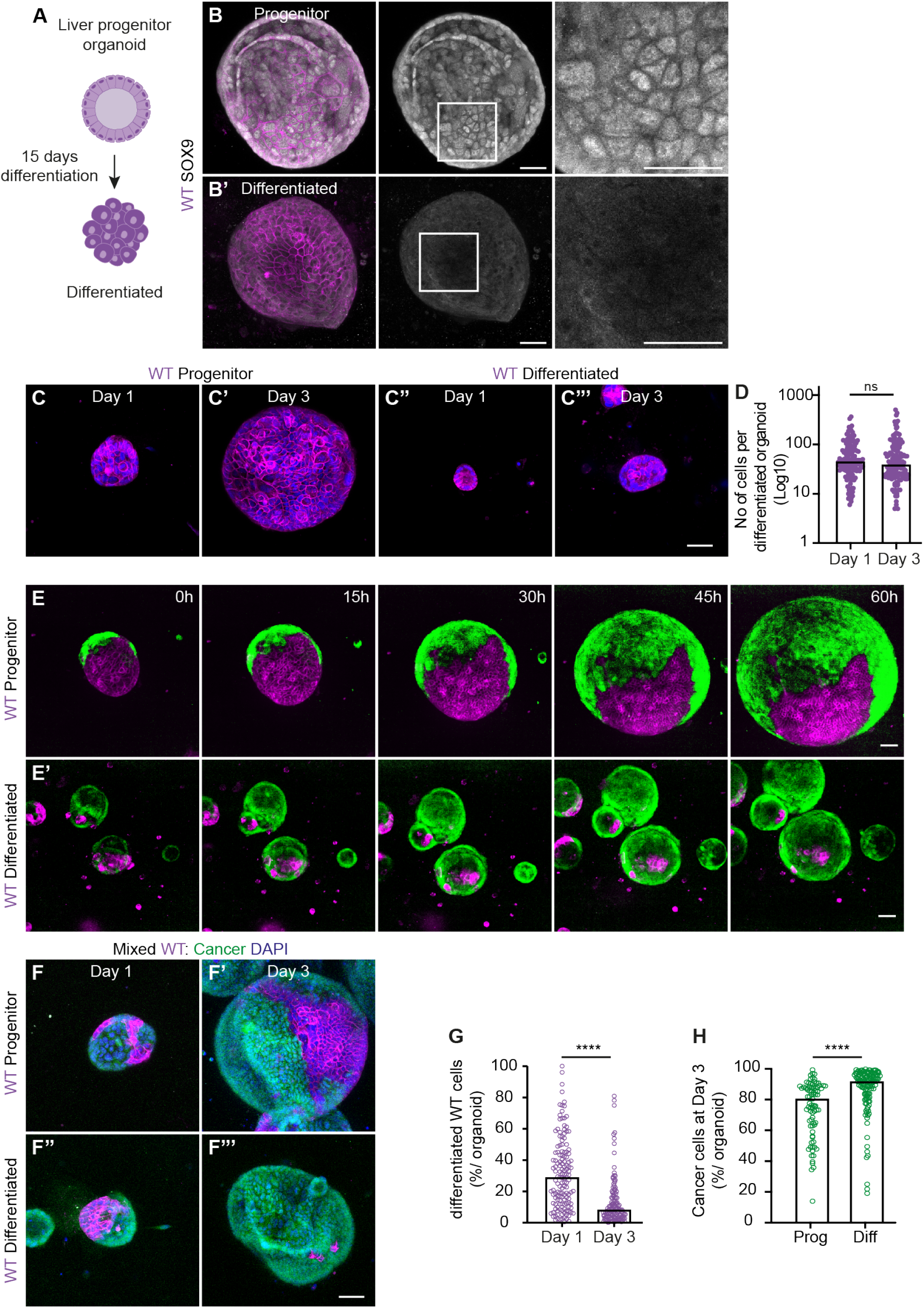
Differentiated WT cells are effectively outcompeted. (A) Schematic depiction of differentiation from progenitor cholangiocytes (top) to hepatocyte-like organoids (bottom) organoids. (B) Representative maximum projections of 3D-confocal images of SOX9 staining in progenitor (B) and differentiated (B’
s) WT organoids. The insets display a 3x magnification of the area in the white box. (C) Representative maximum projections of 3D-confocal images of progenitor (C and C’) and differentiated (C’’ and C’’’) WT organoids, fixed 1 day (C and C’’) and 3 days (C’ and C’’’) after plating; nuclei are visualized with DAPI (blue). (D) Quantification of the number of differentiated WT cells; each dot represents one organoid (Median; Kruskal-Wallis test, Dunn’s multiple comparison test; p>0.9999; n=132 and n=129). (E) Representative maximum projections of 3D-confocal images of time-lapse series of mixed organoids generated from WT progenitor (E) and WT differentiated (E’) cells. (F) Representative maximum projections of 3D-confocal images of mixed organoids from WT progenitor (F and F’) and WT differentiated (F’’ and F’’’) organoids, fixed at day 1 (F and F’’) and day 3 (F’ and F’’’) after plating; nuclei are visualized with DAPI (blue). (G) shows the percentage of differentiated WT cells in mixed organoids at day 1 and day 3. (H) displays the percentage of cancer cells in mixed organoids generated from WT progenitor (left) or differentiated (right) cells; each dot represents one organoid (Median; Kruskal-Wallis test, Dunn’s multiple comparison test; p<0.0001; n=140 and n=128, G; n=81 and n=128, H). Scale bars represent 50μm. See also Figure S5 and Videos S4-S6.

Next, differentiated wild-type organoid cultures were disrupted and aggregated with clumps of cancer cells to generate mixed organoids (Figure S5H). Using time-lapse microscopy, we observed a rapid takeover of the mixed organoids by the cancer population (Figure 5E and Video S6). This effect was obvious from early stages of the experiment, while the effects of competition on progenitor cells first start to appear after 30 hours (Figures 5E, 1F and Video S6). Together, this indicates that outcompetition of differentiated wild-type cells is much more efficient than that of progenitor cells. Indeed, the number of differentiated wild-type cells that contribute to mixed organoids reduced 3.5 times over the course of two days (Figures 5F and 5G) compared to a two-fold reduction of competing progenitor cells (Figures 5F and S5I). This led to mixed organoids that contained over 90% cancer cells three days after plating (Figures 5F and 5H). Thus, differentiated wild-type liver cells are non-proliferative and therefore more susceptible to outcompetition by intestinal cancer cells.

### WT liver acts as a scaffold for tumor growth during competition

Next, we aimed to develop a model which better resembles the overall organ structure of the liver and allows us to study liver-cancer interactions in a self-organizing matrix-free environment. We were inspired by reported microtissues from organs such as brain^45^, heart^46^ and liver^47–50^ that were previously used to study the tissue development and response to injury. For this, pieces of hepatocyte-like differentiated organoids were aggregated in a U-bottom plate to obtain self-organizing liver microtissues with a size of 1-2mm (Figure 6A). To avoid cues coming from the extra-cellular matrix we withdrew the culture matrix during aggregation. These liver microtissues were devoid of SOX9 expression illustrating that the differentiation status of hepatocyte-like wild-type liver cells was conserved (Figure 6B).

**Figure 6:**
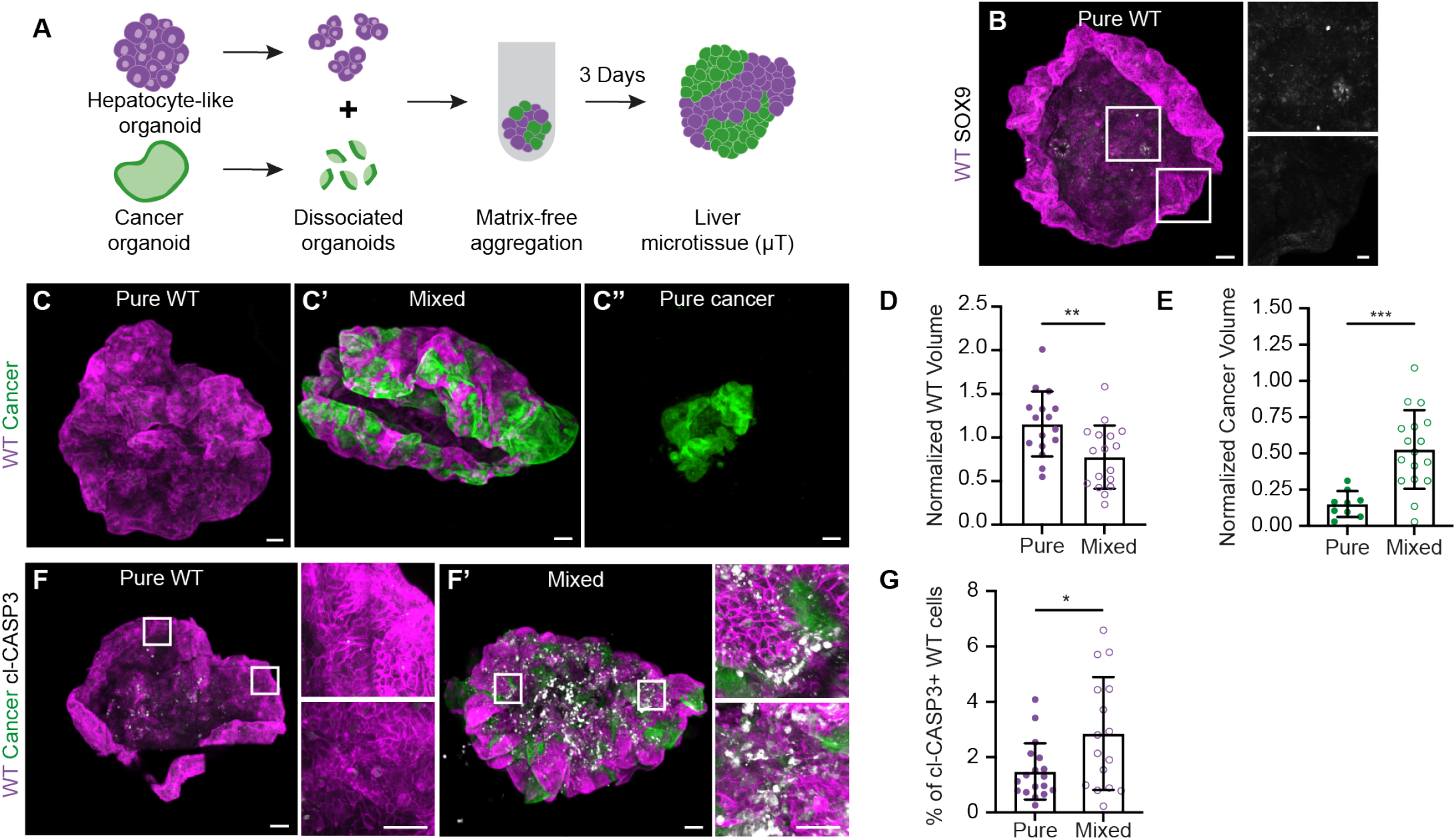
WT liver acts as a scaffold for tumor growth during competition. (A) Schematic depiction of generation of liver metastasis microtissues. (B) Representative 3D-reconstructed stitched confocal images of a pure WT microtissue stained for SOX9 (grey); nuclei are visualized with DAPI (blue). The insets display a 2.5x magnification of the area in the white box. (C) Representative 3D-reconstructed stitched confocal images of from pure WT (C), mixed (C’) and pure cancer (C’’) microtissues. (D-E) Quantification of the WT (D) and cancer (E) volume contributing to pure and mixed microtissues; each dot represents one microtissue (mean ± SEM; unpaired t-test, two-tailed; p<0.0057; n= 16 and 17 microtissues, D; p=0.005; n= 9 and 17 microtissues, E). (F) Representative 3D-reconstructed stitched confocal images of WT pure (F) and mixed (F’) microtissues stained for Cleaved Caspase3 (grey); nuclei are visualized with DAPI (blue). The insets display a 7x magnification of the area in the white box. (G) Quantification of Cleaved Caspase3 positive WT cells within the epithelium of the pure and mixed microtissues; each dot represents one microtissue (mean ± SEM, Mann-Whitney test, two-tailed; p=0.0463; n= 18 and 16 organoids). Scale bars represent 100 μm excluding magnifications in, where scale bar represents 10 μm. See also Figure S6.

To study competitive interactions, we added pieces of cancer organoids during aggregation (Figure 6A). This resulted in liver metastasis microtissues where patches of cancer cells are surrounded by liver tissue (Figure 6C). While characterizing these liver metastasis microtissues we made two observations: 1) pure cancer cells had difficulties to independently form coherent microtissues and had a size that was markedly smaller than that of pure wild-type microtissues (Figures 6C-E and S6). However, when surrounded by liver microtissue, cancer cells thrived (Figures 6C, 6E and S6). 2) In contrast, generation of pure wild-type microtissues was very efficient (Figure 6C and S6), while the volume of wild-type was much reduced in liver metastasis microtissues (Figures 6C, 6D and S6). This indicates that cancer tissue benefits from interactions with wild-type tissue and uses this as a scaffold for expansion. Furthermore, wild-type tissue suffers from the presence of cancer cells.

To understand the nature of the reduction in wild-type liver tissue volume we questioned whether this was caused by induction of cell death. Staining for cleaved-Caspase3 showed a higher number of apoptotic wild-type cells in mixed compared to pure microtissues (Figures 6F and 6G). Interestingly, in these liver metastasis microtissues, cancer and liver tissue intermingled and closely interacted. We observed that cell death was particularly prominent at the interface between the different cell populations (Figure 6F), suggesting that short-ranged interactions are important for elimination of wild-type cells. Thus, we developed a novel matrix-free model to study competition between liver and cancer tissue. Using these liver metastasis microtissues we show that wild-type liver tissue acts as a scaffold for intestinal cancer cells. This subsequently induces elimination of the wild-type cells, which further promotes colonization of liver tissue by cancer cells.

## Discussion

Colonization of secondary organs is one of the most rate-limiting steps in the formation of metastasis and most cancer cells are eliminated by defense mechanisms of the host tissue before metastasis can form.^51^ Upon arrival, cancer cells need to adapt to a novel microenvironment. This requires an intricate interplay between the metastatic cancer cells and a large variety of cell populations in the new organ. A large body of work has focused on interactions of cancer cells with non-epithelial cell populations. In the liver, for example, growth of colorectal cancer metastases is repressed by activated B cells.^52^ In contrast, high levels of liver fibrosis correlate with a poor prognosis for patients with colorectal cancer, indicating a pro-metastatic effect of fibroblasts.^53^ In addition, hepatic stellate cells, liver-specific mesenchymal cells, promotes the exit from dormancy of metastasized breast cancer through induction of a fibrotic injury response.^54^ Even though hepatocytes can assist liver metastasis through a systemic response that facilitates formation of a pro-metastatic niche^55^, direct interactions of metastatic cancer cells with the epithelial cells in the liver are less well-understood. Here, we identified multiple cellular interactions of intestinal cancer with the two main epithelial cell types of the liver. This multi-step process benefits the growth of cancer cells through outcompetition of healthy wild-type cells. Interestingly, such cellular interactions have previously been suggested based on histopathological growth patterns of liver metastasis.^15^ Especially, the replacement pattern, which shows direct interactions and intermingling of liver tissue and cancer cells and is correlated with a poor outcome for patients. However, these studies are primarily based on end-point analysis of histopathological samples or imaging methods that do not reach cellular resolution. The culture models that we have developed here allow the dynamic analysis of such competitive interactions with a subcellular and temporal resolution. Importantly, each dedicated model is optimized to study aspects of this multi-step competition process.

We previously showed that, at the primary site, competition driven by intestinal cancer cells can enforce a cell state transition of surrounding healthy small intestinal cells.^27^ Interestingly, an analogous response is provoked in healthy liver tissue by the same intestinal cancer cells. However, during competition at the primary site, the neighboring intestinal wild-type cells dedifferentiate, while instead at the metastatic site, progenitor cells undergo cell cycle arrest and differentiation. Remarkably, in both tissues this is the natural response to tissue damage; In the intestine, injury causes reversion to a primitive fetal-like state^29,30,56^ and chronic injury in the adult liver induces differentiation of cholangiocytes into hepatocytes through the so-called ductal reaction.^9,10^ Together, this suggests that surrounding organ tissue reacts to cancer-driven competition by activation of an injury response that is embedded in the host tissue. In both tissues, activation of an injury response and the subsequent outcome of competition are similar and result in elimination of the wild-type population and increased growth of the cancer population. However, the steps that are taken to reach this outcome are tissue-specific. Where intestinal cells are rapidly forced to undergo apoptosis, the initial effect of competition on wild-type liver progenitors is compaction and initiation of a cell cycle arrest. Only differentiated cells are eventually eliminated via apoptosis. This has a major effect on the timing of events and the impact on the cancer population. Where inhibition of apoptosis is sufficient to block competition of primary intestinal cancer, this will unlikely prevent loss of liver tissue. In fact, the cancer population initially benefits from the presence of wild-type tissue. For example, competitive interactions with progenitor cells causes increased expansion of cancer cells and interactions with differentiated cells provide a scaffold for cancer cell colonization. Understanding how this feedback is regulated, in terms of the supply of niche factors, identification of involved paracrine signaling pathways or mechanical interactions is an important aim for further research.

The here described liver metastasis microtissues were inspired by microtissues from different organs such as brain^45^ or heart^46^, which have been instrumental to understand the function and response of the tissue of origin to different types of damage or drug response. Here, they were crucial to identify competitive interactions between differentiated liver tissue and cancer cells. In particular, the requirement of the wild-type tissue to provide a scaffold for colonization by cancer cells. Previously, liver microtissues with a functional biliary ductal network were generated, through aggregation of hepatocytes, cholangiocytes and fibroblasts^57^. Combining these cell types with intestinal cancer cells would allow full recapitalization of the major cellular interactions during liver metastasis. Importantly, the exclusion of immune cells from this culture system allows us to untangle the complex cellular interplay without interference of the immune system. It would open the possibility to model other histopathological growth patterns of liver metastasis^15^, such as the desmoplastic type, which is characterized by interactions with stromal cells. In addition, the models are easily adaptable to other cancers that are prone to metastasize to liver, such as breast and pancreatic cancer. Together, our findings can be used to create a true molecular understanding of how healthy cells and cancer cells interact and influence each other, which is a critical step forward towards competition-based cancer therapy.

## Acknowledgments

We thank Ilya Grigoriev and the Biology Imaging Center of Utrecht University Centre for technical support with imaging; members of the Division of Developmental Biology for input and technical support. Jacco van Rheenen for critically reading the manuscript. This work was financially supported by Dutch Cancer Society Young Investigator Grant 11491 / 2018-1 (to S.J.E.S.) and Ayudas Margarita Salas para la formación de jóvenes doctores CA4/RSUE/2022-00236 (to M.L.T.).

## Methods

### Culture of mouse organoids

Please refer to the tables (below) for the composition and recipes of all culture media and solution. Wild-type cholangiocytes were derived from Rosa26-CreERT2::mTmG^58^ females 8-21 weeks, FUCCI2^36^ males 8-21 weeks, Rosa26-rRtTA-H2BmCherry males 8-21 weeks as previously described^32^. In short, isolated liver tissue was minced on a petri dish and incubated in a digestion solution (Collagenase A and Dispase II). After digestion, several rounds of centrifugation were used to discard debris and isolate ducts. Isolated ducts were plated in Cultrex PathClear Reduced Growth Factor Basement Membrane Extract Type 2 (BME2) in mouse isolation medium. Small intestine cancer organoids, derived from the small intestine of Villin-Cre^ERT2^Apc^fl/fl^Kras^G12D/WT^Tr53^fl/R172H^ mice were previously reported.^33^ All lines were cultured as described previously^31^ in drops of BME2 in mouse liver expansion or isolation medium + Noggin.

### Transduction of organoids

Lentiviral transduction was preformed using standard procedures. In short, lentivirus was produced in HEK293T by co-transfection of a dual lentiviral vector 3^rd^ generation pLentiPGK Hygro DEST H2B-mCerulean3 with helper plasmids pMDLg/pRRE, pRSV-Rev and pMD2.G (gifts from Markus Covert and Didier Trono, Addgene plasmids #90234, #12251, #12253 and #12259). Viral particles were harvested from cells four days after transfection and concentrated using 50kDa Amicon Ultra-15 Centrifugal Filter Units (Merck, cat#UFC905024). Organoids were dissociated by mechanical disruption and dissolved in 250 μL ENR medium supplemented with Y-27632 and Polybrene together with the concentrated virus. Cells were incubated at 32°C while spinning at 600xG for 1 hour followed by a 4-hour incubation at 37°C before plating in BME2. Selection was carried out from day 3 onwards with hygromycin.

### Generation of mixed organoids

Mixed organoids were generated as described previously.^31^ In short, suspensions of small clumps of cells were generated from organoids by mechanical disruption and divided over Eppendorf vials in a 1:1 ratio (WT: cancer). Cells were concentrated by mild centrifugation and the pellet was resuspended in a small volume of mouse liver expansion medium and incubated at 37°C for 30 minutes. Cell aggregates were plated in BME2 and cultured in mouse liver expansion medium. For imaging purposes cells were plated in μ-Plate 96 well black uncoated plates or 15μ-Slide 8 well^high^ uncoated plates.

### Differentiation

Cholangiocyte organoids were subjected to a 15-day differentiation protocol developed by STEM CELL Technologies (“Initiation, growth and differentiation of human organoids using HepatiCult™”) following the suppliers’ guidelines. In short, organoids were maintained in expansion medium for 5 days with a medium refreshment at day 3. On day 5 differentiation medium was added and refreshed every 3 days (on day 8, 11 and 14). At day 15 differentiated organoids were ready for use. For generation of mixed organoids the cultures were supplemented with Noggin at day 14. Cancer organoids were cultured in with differentiation medium + Noggin for at one day prior to mixing.

### Generation of microtissues

Cancer organoids and hepatocyte-like organoids obtained by differentiation (described above) were kept in differentiation medium + Noggin for 24h prior to microtissue formation. Organoids were harvested and wash three times with basic medium to remove the culture matrix. After the last wash, the organoids were mechanically disrupted pipetting through a fire-polished glass Pasteur pipet. Cancer organoids were intensively disrupted to generate small pieces and wild-type organoids were treated with a gentle disruption to prevent induction of an injury response. The disrupted organoids were divided over Eppendorf vials in a 2:1 ratio (WT liver : cancer) and concentrated by gentle centrifugation. Cell pellets were dissolved in 5-8 μL differentiation medium +Noggin and incubated at 37°C for 30min. After aggregation, 100 μL /well of differentiation medium +Noggin was added and divided over 96w U bottom plates. Microtissues were allowed to mature at 37°C for 3 days.

### Immuno-fluorescence

Immuno-fluorescence assays were performed as previously described^31^, while protected them from light. In short, organoids were fixed with 4% Paraformaldehyde (PFA) in PBS for 20-30 minutes. Microtissues were fixed by removal of 50 μL medium and addition of 50 μL of 8% PFA in PBS for 20min. Fixation was followed by a minimum of three washes with PBS0 and samples were stored in PBS0 at 4 °C until use. Samples were permeabilizated and blocked in PBS/0.5% BSA/ 0.5% TX-100 for at least 30 minutes and incubated with primary antibodies overnight at 4 °C. After washing three times in PBS/ 0.1% TX-100 samples were incubated with secondary antibodies and DAPI for from 1-3 hours at RT (organoids) up to overnight at 4 °C (organoids and microtissues)^5^. After three washes with PBS/ 0.1% TX-100 and twice with PBS the samples were kept in PBS for imaging. Microtissues, were mounted in RapiClear clearing solution on glass slides using iSpacers. For EdU detection, organoids were treated with EdU staining solution for 30 minutes prior to antibody incubation.

### Microscopy

Imaging of fixed samples and time-lapse videos of FUCCI2 were acquired on a Carl Zeiss LSM880 Fast AiryScan Confocal Laser Scanning microscope (Axio Observer 7 SP with Definite Focus 2) equipped with a CO2 and 37 °C incubator. A Plan-Apochromat 20x/0.8 WD=0.55mm air objective was used to obtain 12bit images with a 1024 resolution, via bidirectional imaging, using a pinhole size of 1AU. A Z-slice thickness of 2.5 μm (fixed samples) and 5 μm (timelapse and microtissues) was used to cover the complete thickness of the sample (17 to 150 slices). For time-lapse imaging a time interval of 2 hours was used for a total duration of 80 hours. The following laser lines were used: 405nm, Laser Argon Multiline (445/488/514), 561nm and 633 nm.

Live-imaging of pure and mixed organoids containing mTmG-labeled wild-type cells were acquired with an Eclipse Ti2-E with PFS (Nikon, Japan) equipped with a Confocal Spinning Disc Unit CSU-W1-T1 (Yokogawa, Japan). A Plan Apo λD 20x / 0.80, WD=0.80, MRD70270 (Nikon, Japan) dry objective was used to obtain 16bit images with a 1024 resolution and 2×2 binning. A Z-slice thickness of 5 μm was used to cover the complete thickness of 300 μm. The time interval was 5 hours for a total duration of 60-70 hours. The following laser lines were used: Stradus 445 (441 nm / 80 mW, Vortran, USA), Stradus 488 (490 nm / 150 mW, Vortran, USA) and OBIS 561 (561 nm, 150 mW, Coherent, USA). Temperature control and CO2: STXG-PLAMX-SETZ21L (TokaiHit, Japan) were set at 37 °C and 5% CO2.

### Image Analysis

Image processing and analysis were performed using the open-source platform FIJI^59^ in combination with computational routines developed in the programming language Julia. For each raw image, program was developed that facilitated the selection of individual organoids in the confocal and lateral planes of the raw image based on the DAPI channel. Next, a general computational pipeline was applied for each organoid; 1) kernel size definition, 2) image processing and 3) 3D object classification and analysis:

#### 1) Kernel size definition

The kernel size, a theoretical value that is equal to the minimal size of the objects of interest inside of the image, was identified by measurement of the raw image after application of a rude processing pipeline, facilitating the automatic high-throughput processing of all the images. The processing pipeline was based on the usage of the average plus the standard deviation as the value to distinguish between foreground and background, and the application of a median filter (with a kernel size equal to the pixel resolution of the image). Lastly, the Euclidean Distance Transform was calculated and used to detect the maximum values. This correlates to the radius of the objects of interest (kernel size) in the 5% of confocal slices with the highest measurements of optical density, which corresponded to planes with the highest biological content. Finally, the kernel size was determined as the average of these values. For time-lapse data (Figures 1, 2 and 5), an additional step was added: harmonization of the kernel size for all the frames of the time-lapse by taking the average value of the kernel size calculated for the individual time frames.

#### 2) Image processing

To extract and enhance the biological signal from the different channels of each image specific routines were applied. DAPI Intensities were homogenized, and background subtraction was performed using an adapted Top-Hat algorithm. This involved subtraction of a copy of the raw image after application of a Gaussian blur, minimum and maximum filters, based on the kernel radius calculated in the previous step. Next, a median filter was applied to preserve the shape of the nuclei. Background and foreground were distinguished by setting a cut-off value based on the average intensity of each pixel below the average intensity of the entire confocal slice histogram. The watershed algorithm in Fiji was used for segmentation of the objects in 2D. To avoid sharp borders and false object detection, two rounds of watershed algorithms were applied, with an intermediate step of median filtering (kernel size equal to the pixel resolution). The raw binary and segmented images were multiplied with the raw data to preserve the original pixel intensities. In order to detect the centroids, dimensions and orientation of each nuclei, the Object Segmentation and Counter Analytical Resources (OSCAR) model was applied using the parameters and measurements previously described.^46,60^ Dendra-2 (cancer) and mTmG (wild-type) were used to classify the different cell populations and therefore processed in parallel. Both channels were processed using a pipeline that included Gaussian blur, maximum, and minimum filters that were applied to each confocal slice. For the mTmG channel, this was followed by subtraction of the Dendra-2 channel. The resulting images were thresholded using the average intensity of each pixel that was below of the average intensity of the whole confocal slice as a cut-off value. In EdU experiments (Figure 3) H2B-mCherry, expressed wild-type cells, was the sole marker that was used for classification of the cell populations. Therefore, a multi-step pipeline of filters was developed to validate correct classification. First, with a minimum filter was applied, followed by an adapted Top-Hat algorithm (kernel radius not multiplied by two), and a median filter with half the value of the kernel radius to smooth object borders. Outliers were removed to eliminate small patches of expression inside nuclei. The image was then binarized and multiplied by the raw data to preserve the original intensities. EdU & pH3 processing was carried out in three steps. First, the previously explained advanced top-hap algorithm was applied to remove background and equalize the intensities of signal across the image. Then, Fiji routine was used to automatically apply the Li’s Minimum Cross Entropy thresholding method based on the iterative version^61^ over the intensity histogram formed by all confocal slices for each image. For EdU this was followed by application of an opening filter over the binary image to remove small artefacts while preserving the objects. For pH3, we applied a grayscale morphological opening and background subtraction based on a cut-off value that was the average of the intensities that were higher than the average intensity for the entire histogram. For Sox9 the adapted top-hat algorithm was used, followed by application of a median filer and outlier detection in order to smoothen the signal.

#### 3) 3D object classification and analysis

The last step of our analysis is used to extract 3D information from the processed images at a single cell level. To achieve this, the DAPI channel was subjected to an artificial intelligence routine that was specifically designed to detect objects in densely packed environments. Our routine has been thoroughly tested and optimized for the phenotypical characterization of 3D nuclei in multicellular samples of embryos and hearth microtissues.^46,60^ The spatial dimensions of each 3D nucleus were used to quantify the volume of positive signal within each nucleus (referred to as marker volume) and calculation of the average intensity. For classification of cells (e.g. wild-type or cancer), the parameter alpha was introduced, which represents the ratio between the marker volume and the total volume. For the detection of wild-type cells using mTmG, EdU, and pH3 markers, an alpha value of 0.1 was used. For classification of wild-type cells based on H2B-mCherry alpha was set to 0.25. Classification of SOX9 expression was based on the relative intensity of SOX9 in all wild-type nuclei. For this, the individual experiments were first normalized based on the maximum intensity in the experiment. Next, the relative intensities of all wild-type nuclei were combined to determine the distribution of SOX9 expression within the whole cell population (See Figure S4). The 25 and 75 percentile values were used as threshold to respectively define SOX9-low and SOX9-high cells.

Imaris software was used for quantification of FUCCI2, compaction and microtissue experiments. Quantification of cell number and volume was performed on 3D reconstructed images. In short, individual nuclei were segmented using the ‘‘spots’’ function and classified as wild-type (Dendra2-) or cancer (Dendra2+). The average distance to the five closest neighbors was calculated for each nucleus as a proxy for internuclear distance. For FUCCI2 quantifications, the average intensity of hCDT1-mCherry and hGeminin-mVenus was measured. Thresholding was used to classify wild-type nuclei as G1 (mCherry+/mVenus-), S-G2-M (mCherry-/mVenus+) or G0-like (mCherry-/mVenus-).

3D reconstructions of microtissues from five independent differentiation experiments (Figure S6) were analyzed. The ‘‘surface’’ function was used to mask mTmG (wild-type) and Dendra2 (cancer) populations in pure and mixed microtissues and the total volume of the surfaces was calculated.

Quantification of Cleaved-Caspase3 was performed on 3D reconstructions of microtissues from four independent differentiations. For this, three crops with a dimension of 250 μm x 250 μm x 50 μm (X/Y/Z) were made in a blinded fashion. All quantifications were performed by a different person to avoid bias. First, wild-type nuclei were segmented and counted using the “spots” function in the DAPI channel. Next, in mixed conditions the “surface” function was used to classify wild-type cells. Lastly, the number of Cleaved-Caspase3+ cells within the wild-type epithelium was determined (excluding debris and extruded cells).

### Visualization

Images were either 3D reconstructed in Imaris or a maximum projection was made in Fiji (indicated in the figure legends). For microtissues, raw images were first stitched using Zen software and colors were adjusted in Imaris prior snapshots were taken Images and movies were converted to RGB using FIJI, cropped and when necessary, smoothened, cropped, rotated and contrasted linearly.

### Statistical analysis

Statistics were performed using GraphPad Prism. Paired or unpaired t test was used when data showed normal distribution (verified with a normality test), whereas Mann-Whitney test was used for data that did not display parametric distribution. For multiple comparisons we used one-way ANOVA for normal distribution data and Kruskal-Wallis test when data was not showing parametric distribution. Adoption of one statistical test or the other is indicated for each experiment in the figure legends.

## Resources

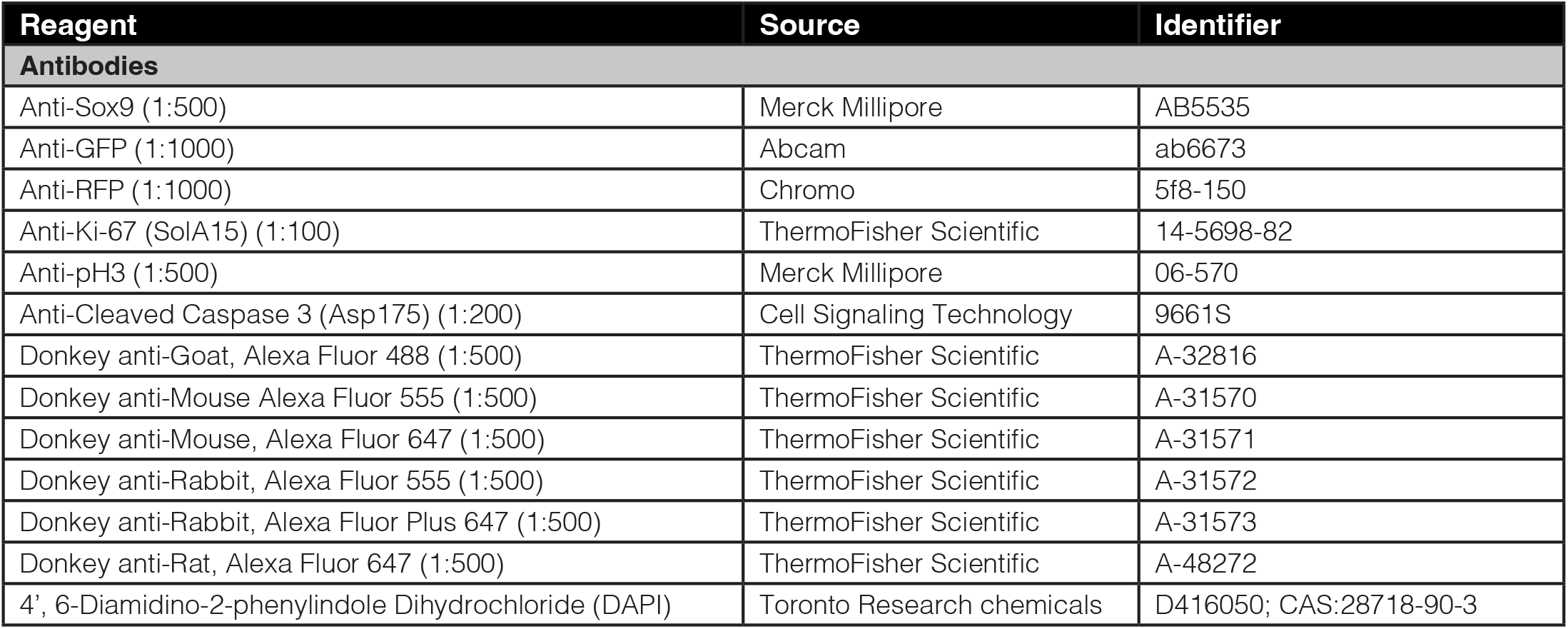
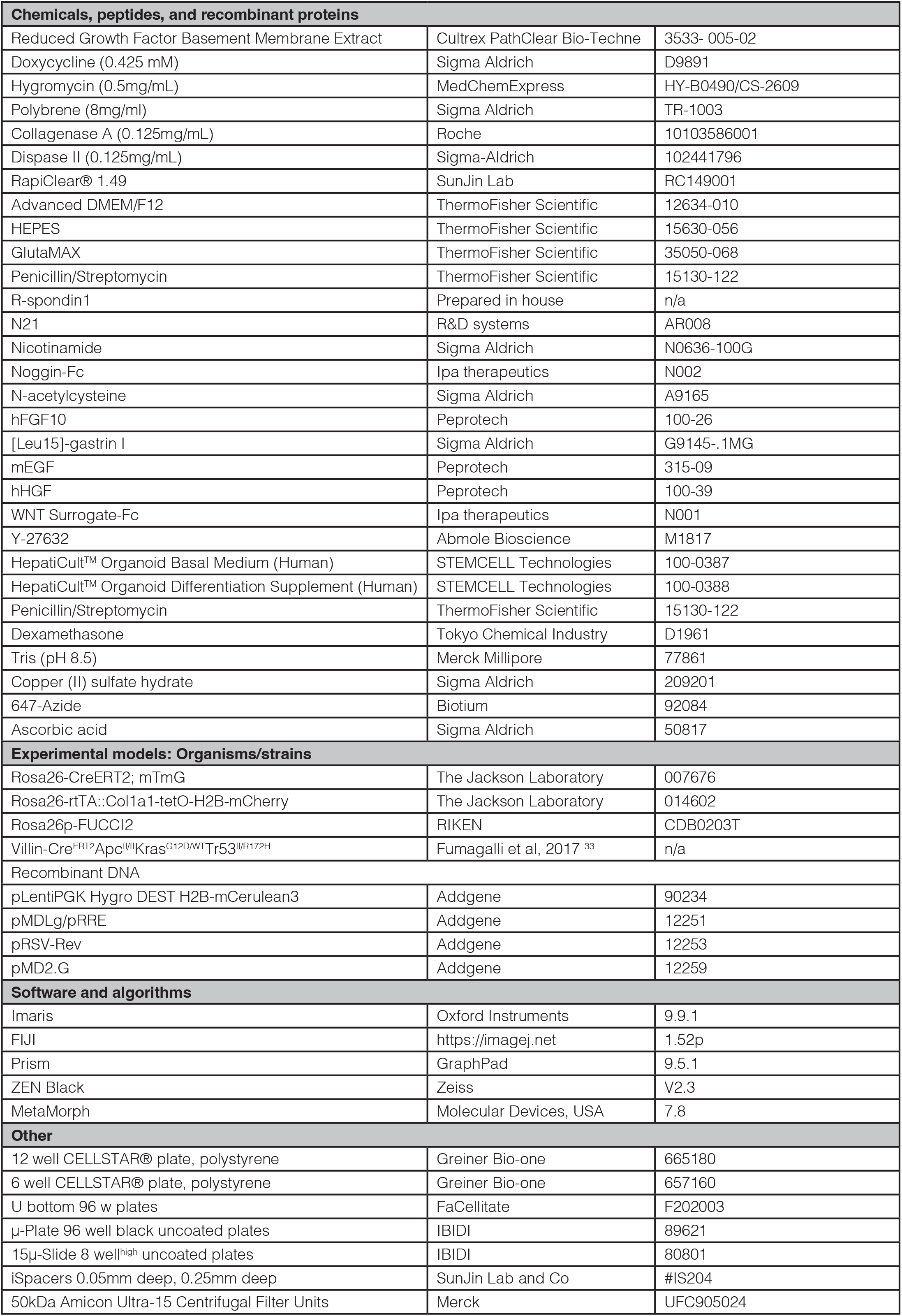

## Solution recipes

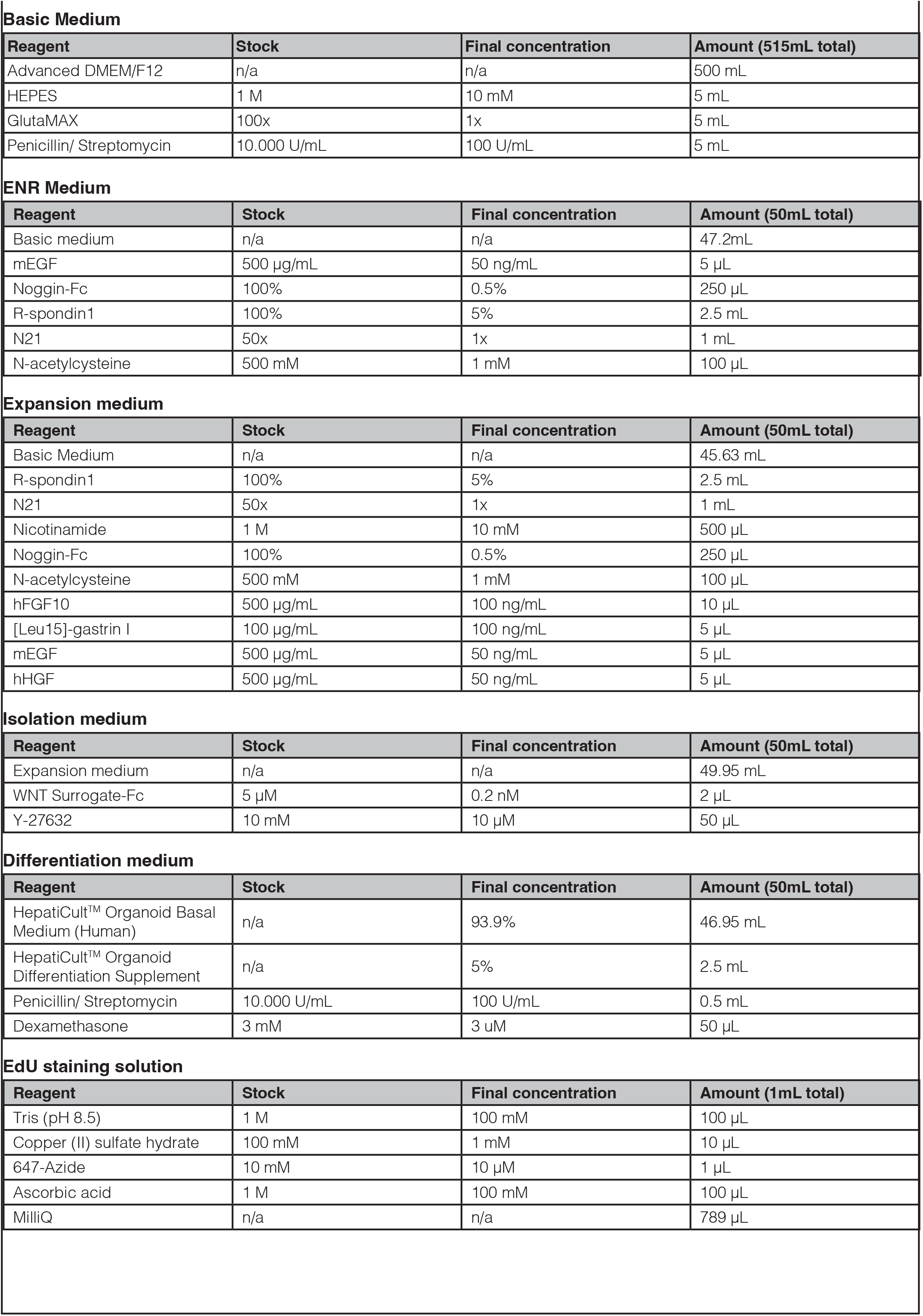

Solution recipes Basic Medium
| Reagent | Stock | Final concentration | Amount (515mL total) |
| --- | --- | --- | --- |
| Advanced DMEM/F12 | n/a | n/a | 500 mL |
| HEPES | 1 M | 10 mM | 5 mL |
| GlutaMAX | 100x | 1x | 5 mL |
| Penicillin/ Streptomycin | 10.000 U/mL | 100 U/mL | 5 mL |

ENR Medium
| Reagent | Stock | Final concentration | Amount (50mL total) |
| --- | --- | --- | --- |
| Basic medium | n/a | n/a | 47.2mL |
| mEGF | 500 µg/mL | 50 ng/mL | 5 µL |
| Noggin-Fc | 100% | 0.5% | 250 µL |
| R-spondin1 | 100% | 5% | 2.5 mL |
| N21 | 50x | 1x | 1 mL |
| N-acetylcysteine | 500 mM | 1 mM | 100 µL |

Expansion medium
| Reagent | Stock | Final concentration | Amount (50mL total) |
| --- | --- | --- | --- |
| Basic Medium | n/a | n/a | 45.63 mL |
| R-spondin1 | 100% | 5% | 2.5 mL |
| N21 | 50x | 1x | 1 mL |
| Nicotinamide | 1 M | 10 mM | 500 µL |
| Noggin-Fc | 100% | 0.5% | 250 µL |
| N-acetylcysteine | 500 mM | 1 mM | 100 µL |
| hFGF10 | 500 µg/mL | 100 ng/mL | 10 µL |
| [Leu15]-gastrin I | 100 µg/mL | 100 ng/mL | 5 µL |
| mEGF | 500 µg/mL | 50 ng/mL | 5 µL |
| hHGF | 500 µg/mL | 50 ng/mL | 5 µL |

Isolation medium
| Reagent | Stock | Final concentration | Amount (50mL total) |
| --- | --- | --- | --- |
| Expansion medium | n/a | n/a | 49.95 mL |
| WNT Surrogate-Fc | 5 µM | 0.2 nM | 2 µL |
| Y-27632 | 10 mM | 10 µM | 50 µL |

Differentiation medium
| Reagent | Stock | Final concentration | Amount (50mL total) |
| --- | --- | --- | --- |
| HepatiCult™ Organoid Basal Medium (Human) | n/a | 93.9% | 46.95 mL |
| HepatiCult™ Organoid Differentiation Supplement | n/a | 5% | 2.5 mL |
| Penicillin/ Streptomycin | 10.000 U/mL | 100 U/mL | 0.5 mL |
| Dexamethasone | 3 mM | 3 uM | 50 µL |

EdU staining solution
| Reagent | Stock | Final concentration | Amount (1mL total) |
| --- | --- | --- | --- |
| Tris (pH 8.5) | 1 M | 100 mM | 100 µL |
| Copper (II) sulfate hydrate | 100 mM | 1 mM | 10 µL |
| 647-Azide | 10 mM | 10 µM | 1 µL |
| Ascorbic acid | 1 M | 100 mM | 100 µL |
| MilliQ | n/a | n/a | 789 µL |

## Supplementary information

**Figure S1:**
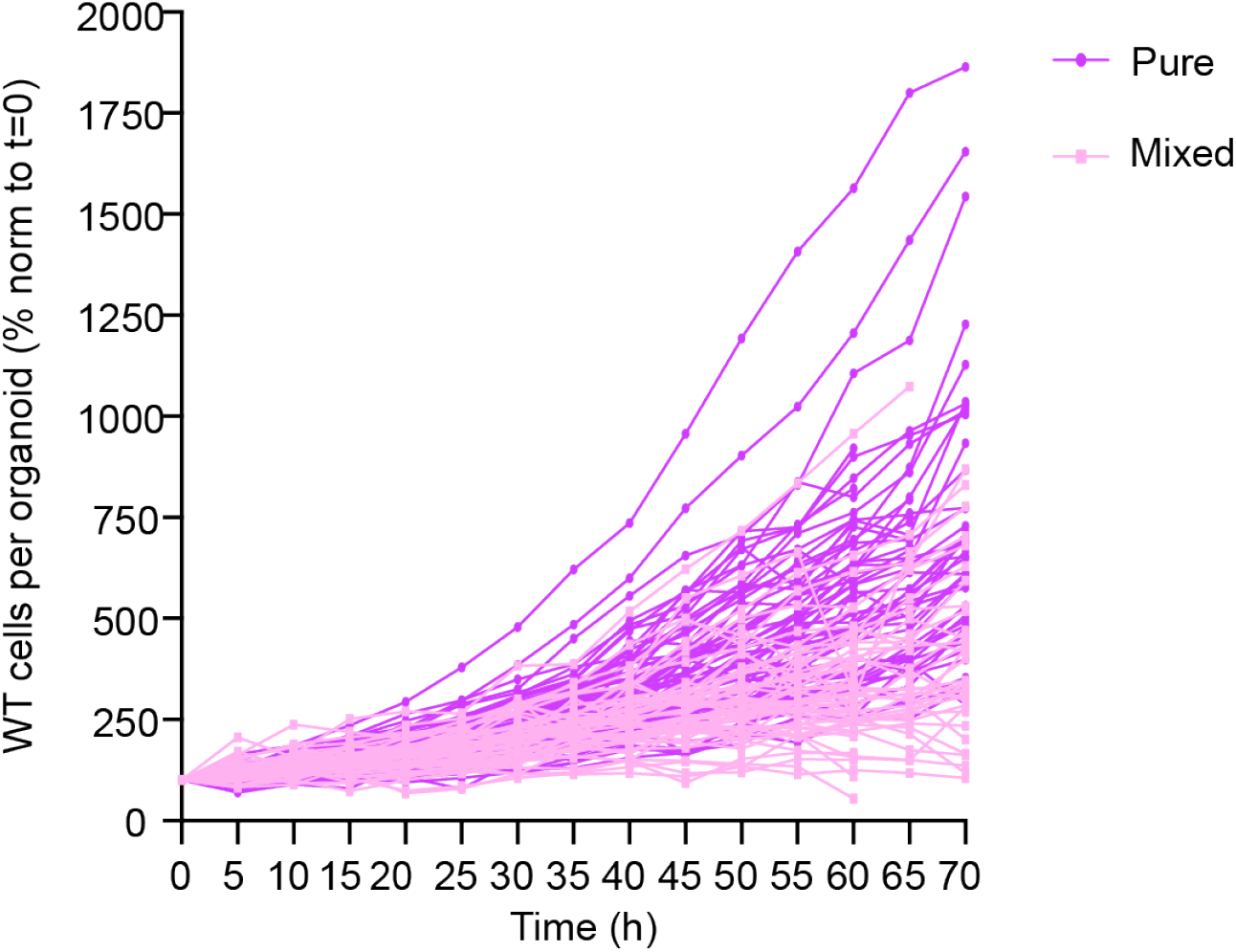
Cancer cells outcompete wild-type liver organoids Related to Figure 1. Displays the wild-type cells in pure (dark purple) and mixed (light purple) that were followed during time-lapse imaging. The number of wild-type cells, normalized to T=0, is plotted against time. Each line represents an organoid.

**Figure S2:**
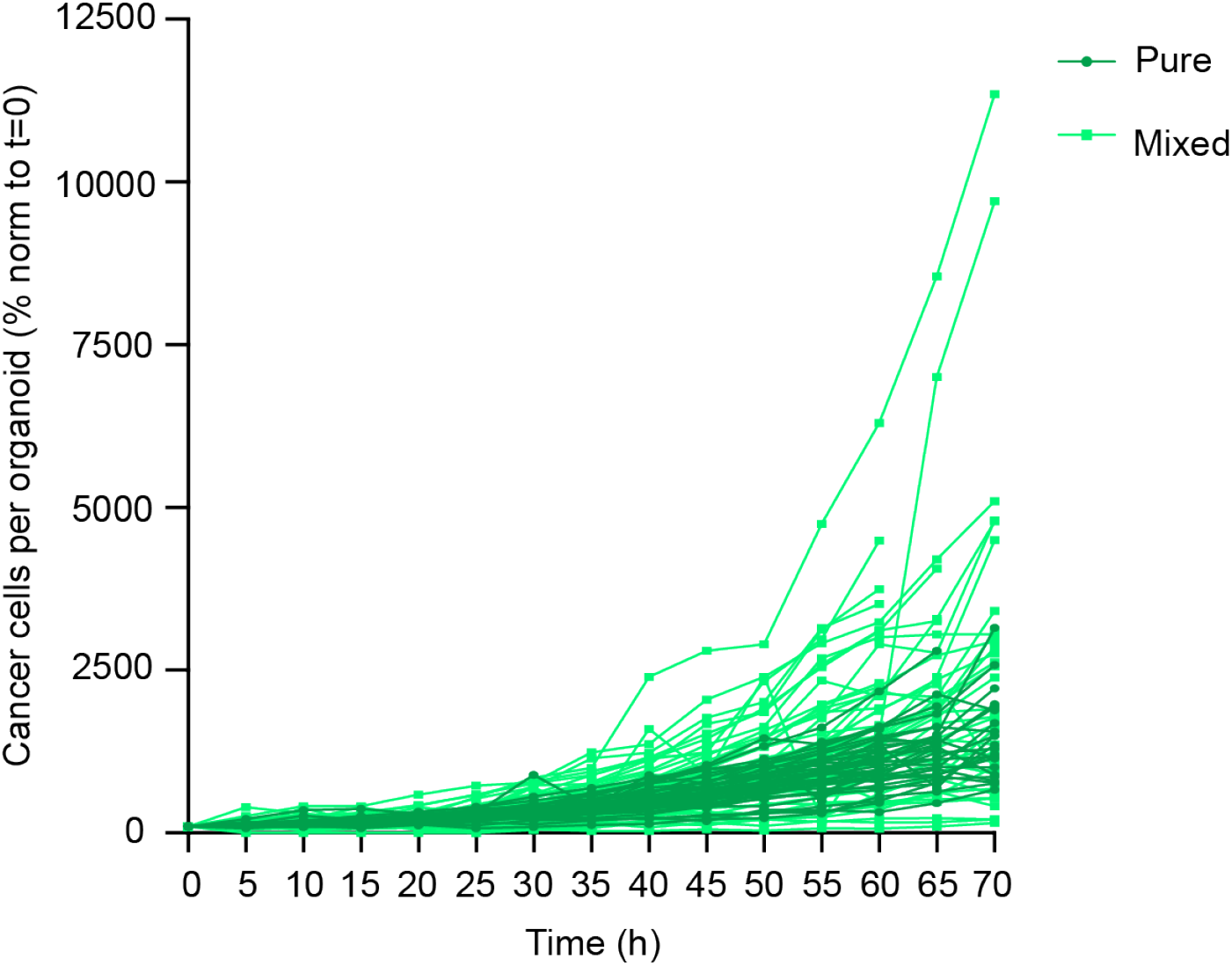
increased expansion of competing cancer cells Related to Figure 2. Displays the cancer cells in pure (dark green) and mixed (light green) that were followed during time-lapse imaging. The number of cancer cells, normalized to T=0, is plotted against time. Each line represents an organoid.

**Figure S3:**
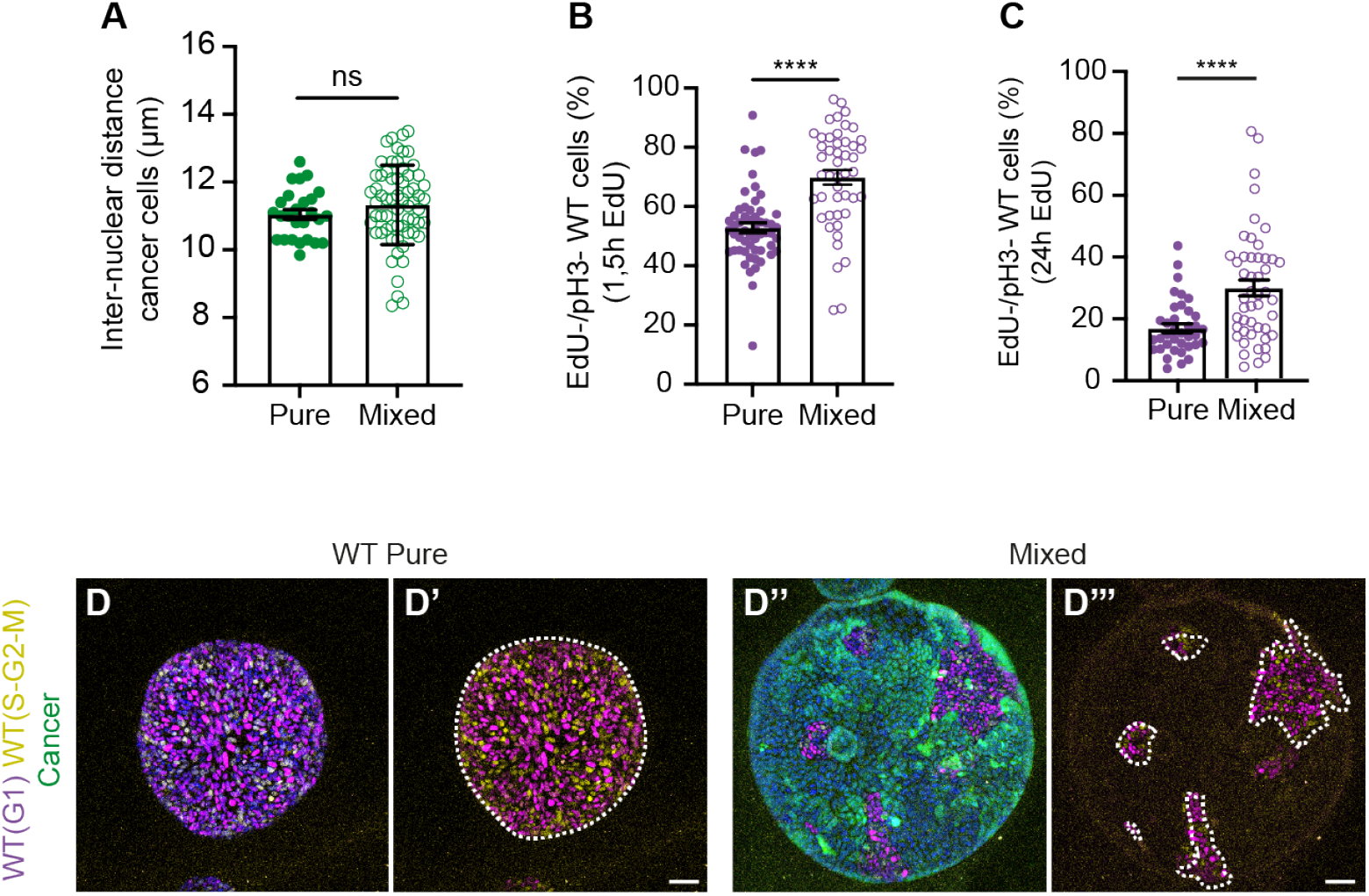
Cancer induces compaction and cell cycle arrest of wild-type liver cells Related to Figure 3. (A) Quantification of the inter-nuclear distance of cancer cells in pure and mixed organoids; each dot represents one organoid (mean ± SEM; Ordinary one-way ANOVA, Sidak’s multiple comparisons test; p<0.4681; n= 26 and 64 organoids). (B-C) Quantification of the percentage of EdU-/ pH3-wild-type cells after 1.5 h (B) or 24 h (C) of EdU treatment; each dot represents one organoid (mean ± SEM; Ordinary one-way ANOVA, Sidak’s multiple comparisons test; p<0.0001; n= 55 and 47 organoids in B; p<0.0001; n=37 and 48 organoids in C). (D) Z projection confocal images of pure wild-type (D and D’) and Mixed (D’’ and D’’’) FUCCI2 organoids fixed three days after planting. White dotted line indicates wild-type cells. Scale bars represent 50 μm.

**Figure S4:**
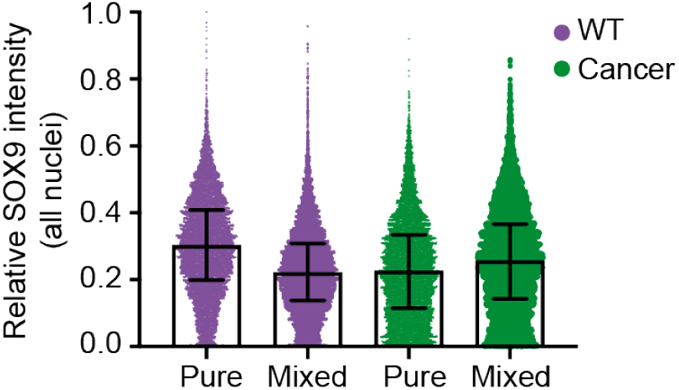
Increased competition through loss of WT progenitor state Related to Figure 4. Displays the relative SOX9 intensities of all individual nuclei in pure and mixed organoids; each dot represents a nucleus (WT pure median =0.2475, n= 7998 nuclei; WT mixed median = 0.1870, n= 9083 nuclei; cancer pure median= 0.2686, n= 8093 nuclei; cancer mixed median= 0.2880, n= 66127).

**Figure S5:**
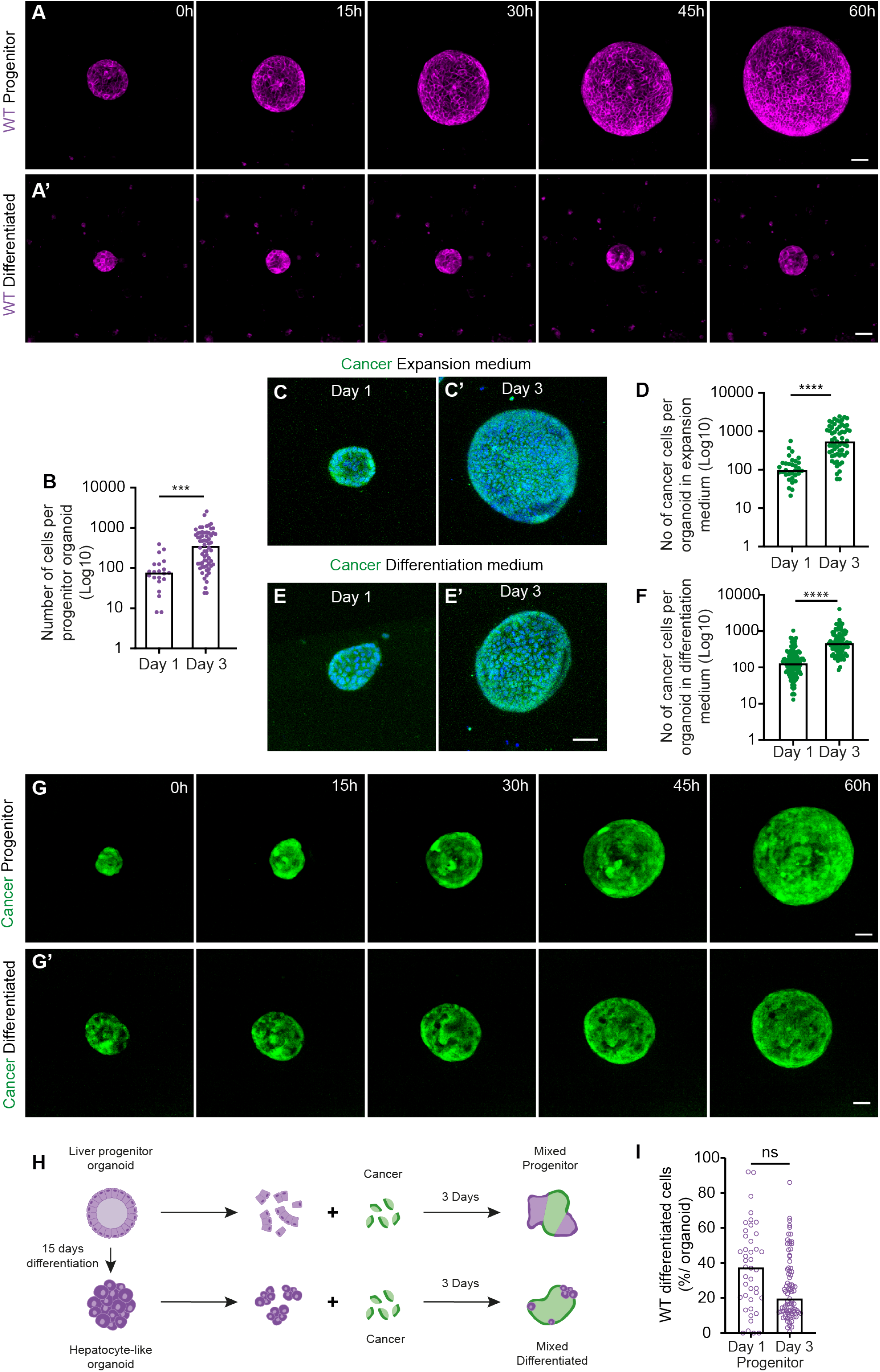
Differentiated WT cells are effectively outcompeted Related to Figure 5. (A) Representative maximum projections of 3D-confocal images of time-lapse series of WT progenitor (A) and WT differentiated (A’) cells. (B) Quantification of the number of WT progenitor cells; each dot represents one organoid (Median; Kruskal-Wallis test, Dunn’
ss multiple comparison test; p=0.0008; n=21 and n=64). (C-F) Representative maximum projections of 3D-confocal images of pure cancer organoids cultured in expansion (C) and differentiation (E) medium, fixed at day 1 (C and E) and day 3 (C’ and E’) after plating; nuclei are visualized with DAPI (blue). Graphs display the number of cancer cells in pure cancer organoids cultured in expansion (D) and differentiation (F) medium; each dot represents one organoid (Median; Kruskal-Wallis test, Dunn’s multiple comparison test; p<0.0001; n=32 and n=61, D; n=124 and n=87, F). (G) Representative maximum projections of 3D-confocal images of time-lapse series of pure cancer organoids cultured in progenitor (G) and differentiation (G’) medium. (H) Schematic depiction of the generation of mixed organoids from progenitor cholangiocyte (top) and differentiated hepatocyte-like (bottom) organoids. (I) shows the percentage of WT progenitor cells in mixed organoids at day 1 and day 3; each dot represents one organoid (Median; Kruskal-Wallis test, Dunn’s multiple comparison test; p=0.1671; n=42 and n=81). Scale bars represent 50 μm.

**Figure S6:**
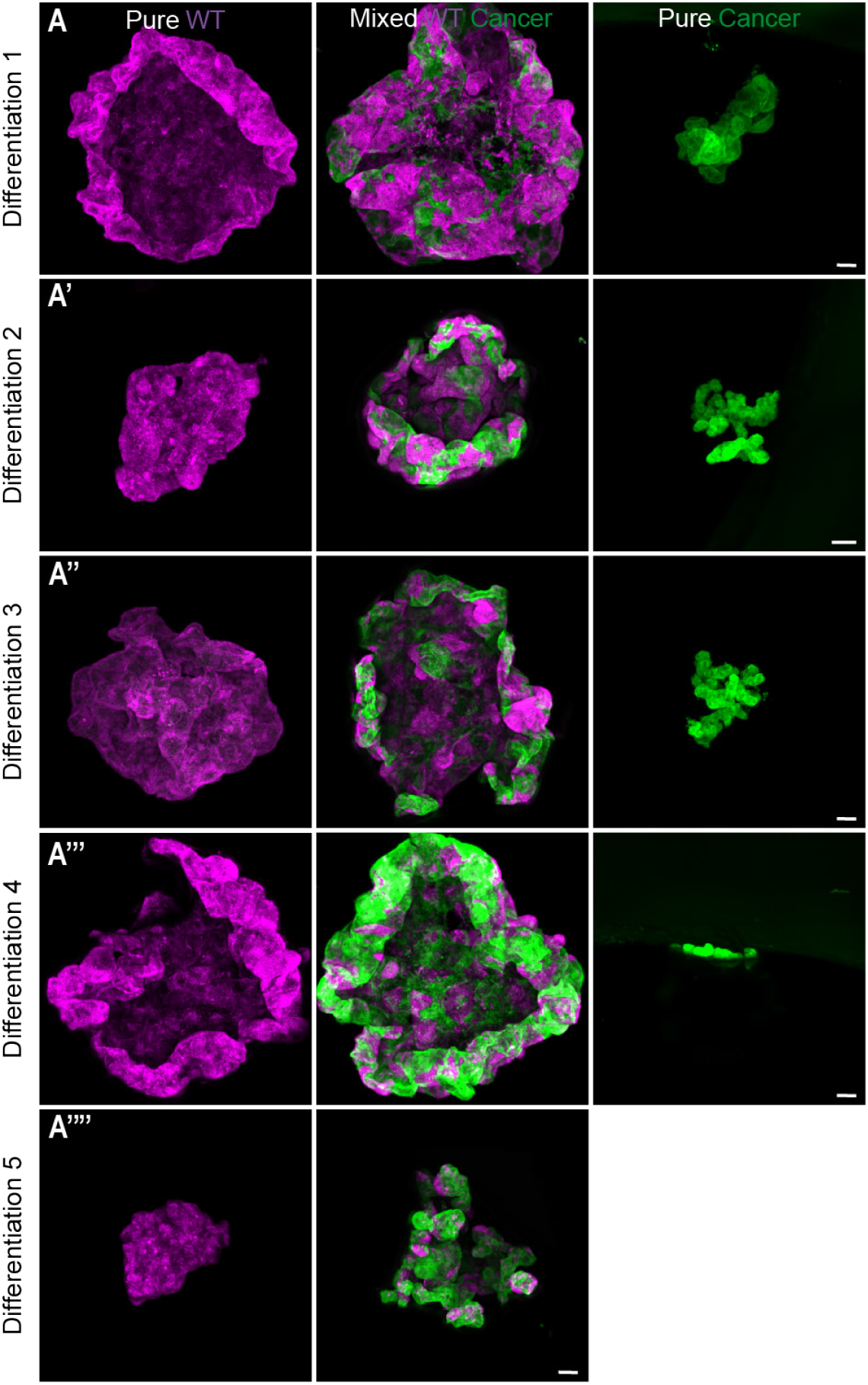
WT liver acts as a scaffold for tumor growth during competition Related to Figure 6. Representative 3D-reconstructed stitched confocal images of from pure WT (left), mixed (middle) and pure cancer (right) microtissues from each round of differentiation. Scale bars represent 100 μm.

## Supplemental Videos

**Video S1: Cancer cells outcompete wild-type liver cells**

Time-lapse series of wild-type (left) and mixed (right) organoids.

**Video S2: Increased expansion of competing cancer cells**

Time-lapse series of cancer (left) and mixed (right) organoids.

**Video S3: Cancer induces compaction and cell cycle arrest of wild-type liver cells**

Time-lapse series of wild-type (left) and mixed (right) FUCCI2 organoids.

**Video S4: No proliferation of differentiated wild-type cells**

Time-lapse series of wild-type progenitor (left) and differentiated (right) organoids.

**Video S5: Differentiation medium does not affect cancer cell expansion**

Time-lapse series of cancer organoids in expansion liver medium (left) and differentiation medium (right).

**Video S6: Differentiated WT cells are effectively outcompeted**

Time-lapse series of mixed organoids with progenitor (left) and differentiated (right) wild-type cells.

## Notes

### Competing Interest Statement

The authors have declared no competing interest.

## References

1. Engstrand, J., Nilsson, H., Strömberg, C., Jonas, E., and Freedman, J. (2018). Colorectal cancer liver metastases – a population-based study on incidence, management and survival. BMC Cancer 18, 78. 10.1186/s12885-017-3925-x.

2. Jones, R.P., Kokudo, N., Folprecht, G., Mise, Y., Unno, M., Malik, H.Z., Fenwick, S.W., and Poston, G.J. (2017). Colorectal Liver Metastases: A Critical Review of State of the Art. Liver Cancer 6, 66–71. 10.1159/000449348.

3. Akgül, Ö., Çetinkaya, E., Ersöz, Ş., and Tez, M. (2014). Role of surgery in colorectal cancer liver metastases. World J Gastroenterol 20, 6113–6122. 10.3748/wjg.v20.i20.6113.

4. Tang, M., Wang, H., Cao, Y., Zeng, Z., Shan, X., and Wang, L. Nomogram for predicting occurrence and prognosis of liver metastasis in colorectal cancer: a population-based study. 10.1007/s00384-020-03722-8/Published.

5. Clark, A.M., Ma, B., Taylor, D.L., Griffith, L., and Wells, A. (2016). Liver metastases: Microenvironments and ex-vivo models. Exp Biol Med 241, 1639–1652. 10.1177/1535370216658144.

6. Mielgo, A., and Schmid, M.C. (2020). Liver tropism in cancer: The hepatic metastatic niche. Cold Spring Harb Perspect Med 10. 10.1101/cshperspect.a037259.

7. Huang, R., Zhang, X., Gracia-Sancho, J., and Xie, W.F. (2022). Liver regeneration: Cellular origin and molecular mechanisms. Liver International 42, 1486–1495. 10.1111/liv.15174.

8. Huch, M., Dorrell, C., Boj, S.F., van Es, J.H., Li, V.S.W., van de Wetering, M., Sato, T., Hamer, K., Sasaki, N., Finegold, M.J., et al. (2013). In vitro expansion of single Lgr5+ liver stem cells induced by Wnt-driven regeneration. Nature 494, 247–250. 10.1038/nature11826.

9. Michalopoulos, G.K. (2007). Liver regeneration. J Cell Physiol 213, 286–300. 10.1002/jcp.21172.

10. Banales, J.M., Huebert, R.C., Karlsen, T., Strazzabosco, M., LaRusso, N.F., and Gores, G.J. (2019). Cholangiocyte pathobiology. Nat Rev Gastroenterol Hepatol 16, 269–281. 10.1038/s41575-019-0125-y.

11. Miyajima, A., Tanaka, M., and Itoh, T. (2014). Stem/Progenitor Cells in Liver Development, Homeostasis, Regeneration, and Reprogramming. Cell Stem Cell 14, 561–574. 10.1016/j.stem.2014.04.010.

12. He, L., Pu, W., Liu, X., Zhang, Z., Han, M., Li, Y., Huang, X., Han, X., Li, Y., Liu, K., et al. (2021). Proliferation tracing reveals regional hepatocyte generation in liver homeostasis and repair. Science (1979) 371. 10.1126/science.abc4346.

13. Malato, Y., Naqvi, S., Schürmann, N., Ng, R., Wang, B., Zape, J., Kay, M.A., Grimm, D., and Willenbring, H. (2011). Fate tracing of mature hepatocytes in mouse liver homeostasis and regeneration. Journal of Clinical Investigation 121, 4850–4860. 10.1172/JCI59261.

14. Miyaoka, Y., Ebato, K., Kato, H., Arakawa, S., Shimizu, S., and Miyajima, A. (2012). Hypertrophy and unconventional cell division of hepatocytes underlie liver regeneration. Curr Biol 22, 1166–1175. 10.1016/j.cub.2012.05.016.

15. Latacz, E., Höppener, D., Bohlok, A., Leduc, S., Tabariès, S., Fernández Moro, C., Lugassy, C., Nyström, H., Bozóky, B., Floris, G., et al. (2022). Histopathological growth patterns of liver metastasis: updated consensus guidelines for pattern scoring, perspectives and recent mechanistic insights. Br J Cancer 127, 988–1013. 10.1038/s41416-022-01859-7.

16. Frentzas, S., Simoneau, E., Bridgeman, V.L., Vermeulen, P.B., Foo, S., Kostaras, E., Nathan, M.R., Wotherspoon, A., Gao, Z., Shi, Y., et al. (2016). Vessel co-option mediates resistance to anti-angiogenic therapy in liver metastases. Nat Med 22, 1294–1302. 10.1038/nm.4197.

17. Moya, I.M., Castaldo, S.A., Van den Mooter, L., Soheily, S., Sansores-Garcia, L., Jacobs, J., Mannaerts, I., Xie, J., Verboven, E., Hillen, H., et al. (2019). Peritumoral activation of the Hippo pathway effectors YAP and TAZ suppresses liver cancer in mice. Science (1979) 366, 1029–1034. 10.1126/science.aaw9886.

18. van Neerven, S.M., and Vermeulen, L. (2023). Cell competition in development, homeostasis and cancer. Nat Rev Mol Cell Biol 24, 221–236. 10.1038/s41580-022-00538-y.

19. Villa del Campo, C., Clavería, C., Sierra, R., and Torres, M. (2014). Cell competition promotes phenotypically silent cardiomyocyte replacement in the mammalian heart. Cell Rep 8, 1741–1751. 10.1016/j.celrep.2014.08.005.

20. Clavería, C., Giovinazzo, G., Sierra, R., and Torres, M. (2013). Myc-driven endogenous cell competition in the early mammalian embryo. Nature 500, 39–44. 10.1038/nature12389.

21. Watanabe, H., Ishibashi, K., Mano, H., Kitamoto, S., Sato, N., Hoshiba, K., Kato, M., Matsuzawa, F., Takeuchi, Y., Shirai, T., et al. (2018). Mutant p53-Expressing Cells Undergo Necroptosis via Cell Competition with the Neighboring Normal Epithelial Cells. Cell Rep 23, 3721–3729. 10.1016/j.celrep.2018.05.081.

22. Kon, S., Ishibashi, K., Katoh, H., Kitamoto, S., Shirai, T., Tanaka, S., Kajita, M., Ishikawa, S., Yamauchi, H., Yako, Y., et al. (2017). Cell competition with normal epithelial cells promotes apical extrusion of transformed cells through metabolic changes. Nat Cell Biol 19, 530–541. 10.1038/ncb3509.

23. Vishwakarma, M., and Piddini, E. (2020). Outcompeting cancer. Nat Rev Cancer 20, 187–198. 10.1038/s41568-019-0231-8.

24. van Neerven, S.M., de Groot, N.E., Nijman, L.E., Scicluna, B.P., van Driel, M.S., Lecca, M.C., Warmerdam, D.O., Kakkar, V., Moreno, L.F., Vieira Braga, F.A., et al. (2021). Apc-mutant cells act as supercompetitors in intestinal tumour initiation. Nature 594, 436–441. 10.1038/s41586-021-03558-4.

25. Flanagan, D.J., Pentinmikko, N., Luopajärvi, K., Willis, N.J., Gilroy, K., Raven, A.P., Mcgarry, L., Englund, J.I., Webb, A.T., Scharaw, S., et al. (2021). NOTUM from Apc-mutant cells biases clonal competition to initiate cancer. Nature 594, 430–435. 10.1038/s41586-021-03525-z.

26. Yum, M.K., Han, S., Fink, J., Wu, S.-H.S., Dabrowska, C., Trendafilova, T., Mustata, R., Chatzeli, L., Azzarelli, R., Pshenichnaya, I., et al. (2021). Tracing oncogene-driven remodelling of the intestinal stem cell niche. Nature 594, 442–447. 10.1038/s41586-021-03605-0.

27. Krotenberg Garcia, A., Fumagalli, A., Le, H.Q., Jackstadt, R., Lannagan, T.R.M., Sansom, O.J., van Rheenen, J., and Suijkerbuijk, S.J.E. (2021). Active elimination of intestinal cells drives oncogenic growth in organoids. Cell Rep 36, 109307. 10.1016/j.celrep.2021.109307.

28. Suijkerbuijk, S.J.E., Kolahgar, G., Kucinski, I., and Piddini, E. (2016). Cell Competition Drives the Growth of Intestinal Adenomas in Drosophila. Current Biology 26, 428–438. 10.1016/j.cub.2015.12.043.

29. Yui, S., Azzolin, L., Maimets, M., Pedersen, M.T., Fordham, R.P., Hansen, S.L., Larsen, H.L., Guiu, J., Alves, M.R.P., Rundsten, C.F., et al. (2018). YAP/TAZ-Dependent Reprogramming of Colonic Epithelium Links ECM Remodeling to Tissue Regeneration. Cell Stem Cell 22, 35–49. e7. 10.1016/j.stem.2017.11.001.

30. Nusse, Y.M., Savage, A.K., Marangoni, P., Rosendahl-Huber, A.K.M., Landman, T.A., de Sauvage, F.J., Locksley, R.M., and Klein, O.D. (2018). Parasitic helminths induce fetal-like reversion in the intestinal stem cell niche. Nature 559, 109–113. 10.1038/s41586-018-0257-1.

31. Krotenberg Garcia, A., van Rheenen, J., and Suijkerbuijk, S.J.E. (2021). Generation of mixed murine organoids to model cellular interactions. STAR Protoc 2, 100997. 10.1016/j.xpro.2021.100997.

32. Broutier, L., Andersson-Rolf, A., Hindley, C.J., Boj, S.F., Clevers, H., Koo, B.-K., and Huch, M. (2016). Culture and establishment of self-renewing human and mouse adult liver and pancreas 3D organoids and their genetic manipulation. Nat Protoc 11, 1724–1743. 10.1038/nprot.2016.097.

33. Fumagalli, A., Drost, J., Suijkerbuijk, S.J.E., Van Boxtel, R., De Ligt, J., Offerhaus, G.J., Begthel, H., Beerling, E., Tan, E.H., Sansom, O.J., et al. (2017). Genetic dissection of colorectal cancer progression by orthotopic transplantation of engineered cancer organoids. Proc Natl Acad Sci U S A 114, E2357–E2364. 10.1073/pnas.1701219114.

34. Wagstaff, L., Goschorska, M., Kozyrska, K., Duclos, G., Kucinski, I., Chessel, A., Hampton-O’Neil, L., Bradshaw, C.R., Allen, G.E., Rawlins, E.L., et al. (2016). Mechanical cell competition kills cells via induction of lethal p53 levels. Nat Commun 7. 10.1038/ncomms11373.

35. Levayer, R., Dupont, C., and Moreno, E. (2016). Tissue Crowding Induces Caspase-Dependent Competition for Space. Current Biology 26, 670–677. 10.1016/j.cub.2015.12.072.

36. Abe, T., Sakaue-Sawano, A., Kiyonari, H., Shioi, G., Inoue, K.I., Horiuchi, T., Nakao, K., Miyawaki, A., Aizawa, S., and Fujimori, T. (2013). Visualization of cell cycle in mouse embryos with Fucci2 reporter directed by Rosa26 promoter. Development (Cambridge) 140, 237–246. 10.1242/dev.084111.

37. Ruijtenberg, S., and van den Heuvel, S. (2016). Coordinating cell proliferation and differentiation: Antagonism between cell cycle regulators and cell type-specific gene expression. Cell Cycle 15, 196–212. 10.1080/15384101.2015.1120925.

38. Kolahgar, G., Suijkerbuijk, S.J.E., Kucinski, I., Poirier, E.Z., Mansour, S., Simons, B.D., and Piddini, E. (2015). Cell Competition Modifies Adult Stem Cell and Tissue Population Dynamics in a JAK-STAT-Dependent Manner. Dev Cell 34, 297–309. 10.1016/j.devcel.2015.06.010.

39. Ellis, S.J., Gomez, N.C., Levorse, J., Mertz, A.F., Ge, Y., and Fuchs, E. (2019). Distinct modes of cell competition shape mammalian tissue morphogenesis. Nature 569, 497–502. 10.1038/s41586-019-1199-y.

40. Liu, N., Matsumura, H., Kato, T., Ichinose, S., Takada, A., Namiki, T., Asakawa, K., Morinaga, H., Mohri, Y., De Arcangelis, A., et al. (2019). Stem cell competition orchestrates skin homeostasis and ageing. Nature 568, 344–350. 10.1038/s41586-019-1085-7.

41. Raven, A., Lu, W.-Y., Man, T.Y., Ferreira-Gonzalez, S., O’Duibhir, E., Dwyer, B.J., Thomson, J.P., Meehan, R.R., Bogorad, R., Koteliansky, V., et al. (2017). Cholangiocytes act as facultative liver stem cells during impaired hepatocyte regeneration. Nature 547, 350–354. 10.1038/nature23015.

42. Antoniou, A., Raynaud, P., Cordi, S., Zong, Y., Tronche, F., Stanger, B.Z., Jacquemin, P., Pierreux, C.E., Clotman, F., and Lemaigre, F.P. (2009). Intrahepatic Bile Ducts Develop According to a New Mode of Tubulogenesis Regulated by the Transcription Factor SOX9. Gastroenterology 136, 2325–2333. 10.1053/j.gastro.2009.02.051.

43. Furuyama, K., Kawaguchi, Y., Akiyama, H., Horiguchi, M., Kodama, S., Kuhara, T., Hosokawa, S., Elbahrawy, A., Soeda, T., Koizumi, M., et al. (2011). Continuous cell supply from a Sox9-expressing progenitor zone in adult liver, exocrine pancreas and intestine. Nat Genet 43, 34–41. 10.1038/ng.722.

44. Kawaguchi, Y. (2013). Sox9 and programming of liver and pancreatic progenitors. Journal of Clinical Investigation 123, 1881–1886. 10.1172/JCI66022.

45. Zamproni, L.N., Mundim, M.T.V.V., and Porcionatto, M.A. (2021). Neurorepair and Regeneration of the Brain: A Decade of Bioscaffolds and Engineered Microtissue. Front Cell Dev Biol 9. 10.3389/fcell.2021.649891.

46. Giacomelli, E., Meraviglia, V., Campostrini, G., Cochrane, A., Cao, X., van Helden, R.W.J., Krotenberg Garcia, A., Mircea, M., Kostidis, S., Davis, R.P., et al. (2020). Human-iPSC-Derived Cardiac Stromal Cells Enhance Maturation in 3D Cardiac Microtissues and Reveal Non-cardiomyocyte Contributions to Heart Disease. Cell Stem Cell 26, 862–879.e11. 10.1016/j.stem.2020.05.004.

47. Hafiz, E.O.A., Bulutoglu, B., Mansy, S.S., Chen, Y., Abu-Taleb, H., Soliman, S.A.M., El-Hindawi, A.A.F., Yarmush, M.L., and Uygun, B.E. (2021). Development of liver microtissues with functional biliary ductular network. Biotechnol Bioeng 118, 17–29. 10.1002/bit.27546.

48. Hong, S., Oh, S.J., Choi, D., Hwang, Y., and Kim, S.-H. (2020). Self-Organized Liver Microtissue on a Bio-Functional Surface: The Role of Human Adipose-Derived Stromal Cells in Hepatic Function. Int J Mol Sci 21, 4605. 10.3390/ijms21134605.

49. Yan, L., Messner, C.J., Tian, M., Gou, X., Suter-Dick, L., and Zhang, X. (2022). Evaluation of dioxin induced transcriptomic responses in a 3D human liver microtissue model. Environ Res 210, 112906. 10.1016/j.envres.2022.112906.

50. Proctor, W.R., Foster, A.J., Vogt, J., Summers, C., Middleton, B., Pilling, M.A., Shienson, D., Kijanska, M., Ströbel, S., Kelm, J.M., et al. (2017). Utility of spherical human liver microtissues for prediction of clinical drug-induced liver injury. Arch Toxicol 91, 2849–2863. 10.1007/s00204-017-2002-1.

51. de Visser, K.E., and Joyce, J.A. (2023). The evolving tumor microenvironment: From cancer initiation to metastatic outgrowth. Cancer Cell 41, 374–403. 10.1016/j.ccell.2023.02.016.

52. Xu, Y., Wei, Z., Feng, M., Zhu, D., Mei, S., Wu, Z., Feng, Q., Chang, W., Ji, M., Liu, C., et al. (2022). Tumor-infiltrated activated B cells suppress liver metastasis of colorectal cancers. Cell Rep 40. 10.1016/j.celrep.2022.111295.

53. Hu, X., Marietta, A., Dai, W., Li, Y., Ma, X., Zhang, L., Cai, S., and Peng, J. (2020). Prediction of hepatic metastasis and relapse in colorectal cancers based on concordance analyses with liver fibrosis scores. Clin Transl Med 9. 10.1186/s40169-020-0264-3.

54. Correia, A.L., Guimaraes, J.C., Auf der Maur, P., De Silva, D., Trefny, M.P., Okamoto, R., Bruno, S., Schmidt, A., Mertz, K., Volkmann, K., et al. (2021). Hepatic stellate cells suppress NK cell-sustained breast cancer dormancy. Nature. 10.1038/s41586-021-03614-z.

55. Lee, J.W., Stone, M.L., Porrett, P.M., Thomas, S.K., Komar, C.A., Li, J.H., Delman, D., Graham, K., Gladney, W.L., Hua, X., et al. (2019). Hepatocytes direct the formation of a pro-metastatic niche in the liver. Nature 567, 249–252. 10.1038/s41586-019-1004-y.

56. Gregorieff, A., Liu, Y., Inanlou, M.R., Khomchuk, Y., and Wrana, J.L. (2015). Yap-dependent reprogramming of Lgr5+ stem cells drives intestinal regeneration and cancer. Nature 526, 715–718. 10.1038/nature15382.

57. Hafiz, E.O.A., Bulutoglu, B., Mansy, S.S., Chen, Y., Abu-Taleb, H., Soliman, S.A.M., El-Hindawi, A.A.F., Yarmush, M.L., and Uygun, B.E. (2021). Development of liver microtissues with functional biliary ductular network. Biotechnol Bioeng 118, 17–29. 10.1002/bit.27546.

58. Muzumdar, M.D., Tasic, B., Miyamichi, K., Li, L., and Luo, L. (2007). A global double-fluorescent Cre reporter mouse. genesis 45, 593–605. 10.1002/dvg.20335.

59. Schindelin, J., Arganda-Carreras, I., Frise, E., Kaynig, V., Longair, M., Pietzsch, T., Preibisch, S., Rueden, C., Saalfeld, S., Schmid, B., et al. (2012). Fiji: An open-source platform for biological-image analysis. Nat Methods 9, 676–682. 10.1038/nmeth.2019.

60. Ledesma-Terrón, M., Pérez-Dones, D., and Míguez, D.G. OSCAR: a framework to identify and quantify cells in densely packed three-dimensional biological samples. 10.1101/2021.06.25.449919.

61. Li, C.H., and Tam, P.K.S. (1998). An iterative algorithm for minimum cross entropy thresholding.

